# A neural signature of pattern separation in the monkey hippocampus

**DOI:** 10.1101/302505

**Authors:** John J. Sakon, Wendy A. Suzuki

## Abstract

The CA3 and dentate gyrus (DG) regions of the hippocampus are considered key for disambiguating sensory inputs from similar experiences in memory, a process termed pattern separation. The neural mechanisms underlying pattern separation, however, have been difficult to compare across species: rodents offer robust recording methods with less human-centric tasks while humans provide complex behavior with less recording potential. To overcome these limitations, we trained monkeys to perform a visual pattern separation task similar to those used in humans while recording activity from single CA3/DG neurons. We find that when animals discriminate recently seen novel images from similar (lure) images, behavior indicative of pattern separation, CA3/DG neurons respond to lure images more like novel than repeat images. Using a population of these neurons, we are able to classify novel, lure, and repeat images from each other using this pattern of firing rates. Notably, one subpopulation of these neurons is more responsible for distinguishing lures and repeats—the key discrimination indicative of pattern separation.

## Introduction

Pattern separation is the putative process the brain uses to differentiate similar inputs into non-overlapping outputs^1^. In the case of memory, when a subject is shown an image similar to one they recently viewed, pattern separation mechanisms are tasked to differentiate the newly presented image from their memory of similar images. Experimental work has found that the CA3 and dentate gyrus (CA3/DG) regions of the hippocampus show neurophysiological signals correlated with successful mnemonic pattern separation in humans^2-6^ and rats^7-11^.

However, there remain key gaps in our understanding of the neural basis of pattern separation from this cross-species literature. For example, human fMRI studies have revealed BOLD responses in the CA3/DG regions of hippocampus characteristic of pattern separation^2-5^ but are limited by spatial and temporal resolution of the BOLD signal. Important studies in rodents have shown electrophysiological correlates of pattern separation of spatial environments in populations of CA3/DG neurons^7-10^, but the differences between the experimental designs used across human and rodent studies make these results difficult to compare. In particular, other than the usual difficulties in comparing BOLD with single neuron responses^12^, human pattern separation tasks typically involve discrimination of visual stimuli from memory on timescales less than 1 second^2-6,13^, while rodent pattern separation tasks involve spatial discrimination of environments over the course of minutes^8-11^. Further, recent findings^14^ raise new questions about the previous electrophysiological signatures of pattern separation in rodents^8-10^.

Here we address this gap between human and rodent pattern separation work by recording from single neurons in the CA3/DG regions of the monkey hippocampus as animals perform a visual pattern separation task closely related to those used in human fMRI work. Tasks that have been used to probe pattern separation fall under two general classes: those that utilize parametric manipulations of repeated stimuli^3,5,15^ or environments^10,16^ and those that use serial presentation of novel, repeat and lure objects^2,4,13^. We elected to utilize the latter paradigm, as the use of novel stimuli in each session engages the animals and avoids confounds associated with long-term memory of stimuli used in previous sessions. Further, the use of an implicit task—unlike tasks where the subject explicitly classifies match from nonmatch stimuli^17^—has been essential in distinguishing signatures of pattern separation in the hippocampus by avoiding conscious recollection^2,5^. Further, explicit classification of stimuli is known to affect responses to images in regions upstream of hippocampus along the ventral visual stream like inferotemporal/perirhinal cortex, where a memory signal develops during delayed match to sample tasks^17^, while this memory signal fails to develop when repeated stimuli are shown over hours or days of passive viewing^18^.

In our task, monkeys serially viewed three categories of images: **Novel** images, which the animal had never seen before; **Lure** images, which had strong visual similarity to a recently seen novel image (examples shown in Fig. S1); and **Repeat** images, which were exact replicas of a recently seen novel image (Fig 1A). Lure and repeat images were shown within 0-3 trials after their respective novel image was shown. Animals were allowed to look at each image as long as desired (up to 10 s) until they ended the trial by saccading outside the bounds of the image. We compiled 19,350 novel images, 6450 repeat images (each a repeat of an image from the novel set) and 6450 lure images (each similar to an image from the novel set), which allowed us to never show the monkeys any novel or lure image more than once or the repeated novel image more than twice.

**Figure 1.**
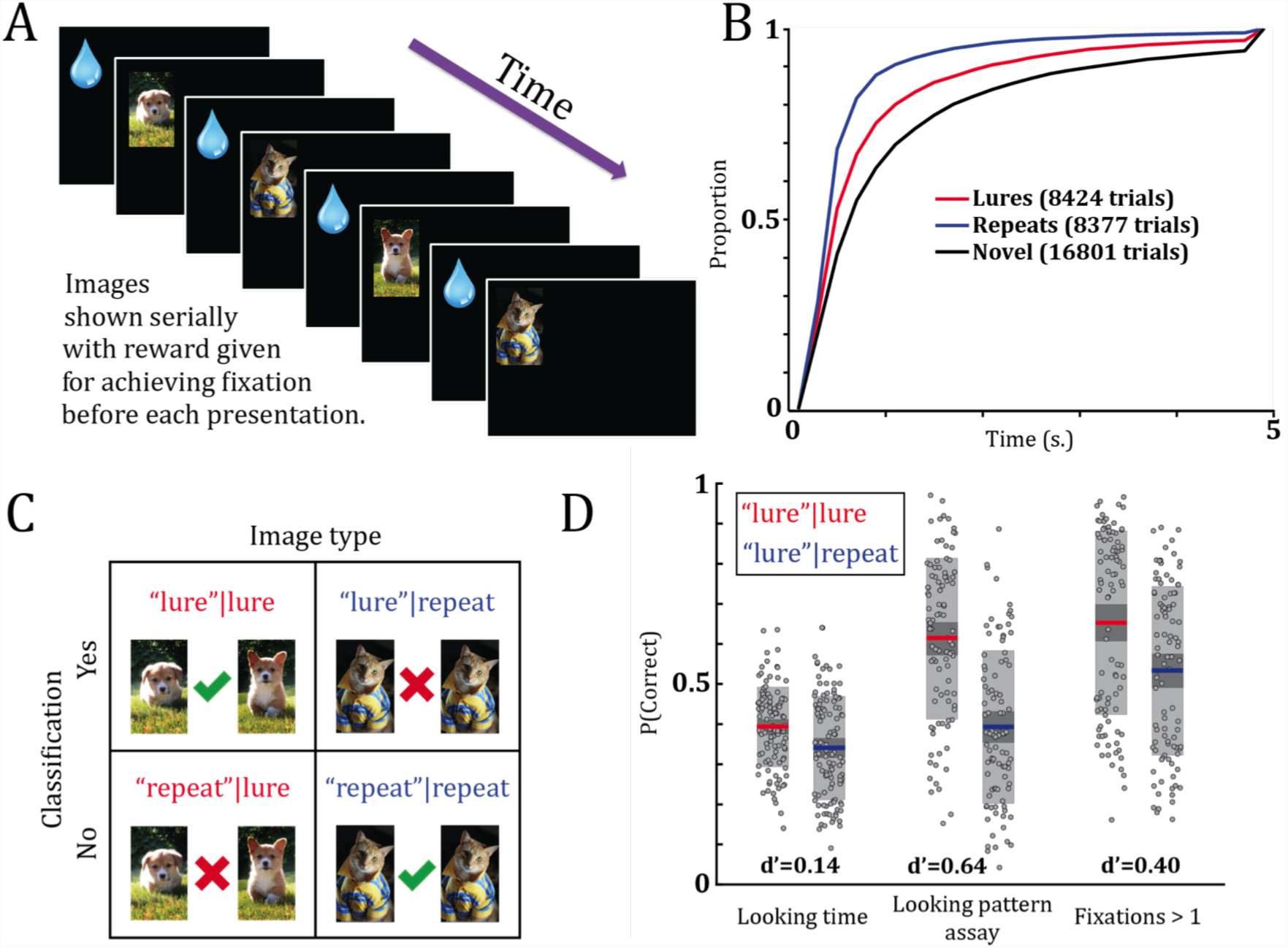
Task outline and behavioral results. **A)** Diagram of task structure. In this example, the monkey would see two novel (never-before-seen) images followed by a lure image (3^rd^ image) and then a repeat image (4^th^ image). Images are centered on the screen. Reward is given before every image presentation when fixation to a center dot is achieved. **B)** Cumulative histogram of looking times pooled across all animals and separated by trial type. Monkeys looked significantly longer at novel compared to repeat images (novel mean=1.40 s. v. repeat mean=0.73 s., p=9.6E-10) and lure compared to repeat images (lure mean=1.04 s. v. repeat mean=0.73 s., p=2.0E-8). Monkeys also looked longer at novel compared to lure images (p=1.1E-8) (Tukey’s HSD of group means after multi-level ANOVA for all pairwise comparisons, Methods), as expected since animals only likely discriminate a proportion of lure images. **C)** Chart of the behavioral classification scheme inspired by signal detection theory. Image type indicates if a lure or repeat was shown to the monkey, while classification indicates how the monkey treated the image based on the behavioral metric. For the looking time analysis, when monkeys looked at a repeat image for shorter than its respective novel image, this was considered a “repeat”|repeat. When monkeys looked at a lure image longer than its respective novel image, we considered it a “lure”|lure. For the looking pattern assay, when **ρ** between a repeat and novel image was higher than threshold, this was considered a “repeat”|repeat. And when **ρ** between a lure and novel image was lower than threshold, we considered this a “lure”|lure. Therefore, when **ρ** is lower than threshold on repeats—a false alarm event—we consider this a “lure”|repeat. And when **ρ** is higher than threshold on lures—a false negative event—we consider this a “repeat”|lure. **D)** Hit rate (a hit being a “lure”|lure) and false alarm rate (a false alarm being a “lure”|repeat) for 88 sessions in 3 monkeys for three classification methods (Methods). Left: looking time assay. Center: looking pattern assay using the best **ρ** threshold value for each session. Right: classification of hits and false alarms using only lures and repeats with >1 fixation. Each method was done with the same threshold for both “lure”|lure v. “repeat”|lure and “lure”|repeat v. “repeat”|repeat comparisons, making the d’ values an unbiased assessment of behavioral classification. Averages are shown by colored lines, shaded boxes are the 95% confidence interval (dark gray) and the standard deviation (light gray).

This implicit task design allows us to address the hypothesis—suggested by human fMRI work on pattern separation^2,3,5^—that CA3/DG neurons treat lure images more like novel images as compared to relatively different responses between repeat and novel images. This prediction is in contrast to neural responses to passive viewing tasks in more perceptual (and less mnemonic) regions along the ventral visual stream such as inferotemporal/perirhinal cortex, where neurons treat repeat images more like novel images by maintaining similar firing rates^18^, as compared to relatively different responses between novel and lure images (with increasing differences in firing rates to less similar images^19,20^). Indeed, utilizing firing rates from our population of CA3/DG neurons, we are able to discriminate novel, lure and repeat images from each other with a pattern of responses similar to that shown in human fMRI studies. Notably, the key lure v. repeat discrimination is only shown by a subpopulation of these neurons, suggesting a signature of pattern separation via hippocampal firing rates.

## Results

### Behavior

We utilized preferential looking to gauge monkeys’ ability to identify repeat and lure images^21,22^. How long monkeys looked at each image before saccading outside each image’s bounds was recorded and pooled across each of our three trial types for 121 sessions in three macaque monkeys (Rhesus BY=55 sessions, Rhesus I=56 sessions, Bonnet B=10 sessions). Consistent with previous studies in monkeys^21^ and infants^22^, we found that animal looking times was significantly less for repeat compared to novel images (Fig. 1B), indicating recognition of previously seen images. We predicted that when monkeys identified a lure image, they would look longer at the lure relative to a repeat image, as the newness of the lure should increase the looking time of a portion of images through dishabituation^22^. Indeed, animals looked significantly longer at lure compared to repeat images (Fig. 1B). Monkeys continued to be engaged in the task across each session, as looking times for lure and repeat images remained consistent throughout (Fig. S2a). Since the variable number of intervening images between each novel/lure and novel/repeat pair (0-3 images) showed no effect on looking time performance (p>0.6, multi-level ANOVA, Fig. S2B-C), all trial types were pooled regardless of number of intervening images.

A possible concern is that, since looking times for lure images are about halfway between looking times for novel and repeat images, the monkeys merely treat the lure images as a mixture of novel and repeat images. In other words, they consider some lure images as novel from the novel images they’ve recently seen, and some lure images as repeated from the novel images they’ve recently seen. This “mixture” hypothesis makes a clear prediction: that the variance of the lure looking times should be equal to the variance of the mixture of novel and repeat looking times^23^. Instead, the variance for lure looking times was significantly smaller than the variance for pooled novel and repeat images (p=9.3E-7, Levene’s test), suggesting that monkeys do not see lure images as merely a mixture of novel and repeat images.

To quantify the monkeys’ ability to discriminate lure and repeat images using preferential looking, we developed a signal detection theory framework with four categories (Fig. 1C). A correctly classified repeat image, where the monkey recognizes the image as a repeat of a recently seen novel image, is called a **“repeat”|repeat** (i.e. true negative, as if the monkey responded “repeat” when shown a repeat), while a correctly classified lure image, where the monkey identifies the lure as different than its previously seen novel partner image, is called a **“lure”|lure** (i.e. true positive). **“lure”|repeat** is a repeat that is incorrectly classified as a lure (i.e. false positive), and **“repeat”|lure** is a lure incorrectly classified as a repeat (i.e. false negative). This analysis achieved a hit rate of 0.39±0.01 and a false alarm rate of 0.34±0.01, which yielded signal detection at d’=0.14 (Fig. 1D).

In addition to looking times, a second signature of monkey behavior is the eye scanpath as the monkeys observe images. We developed a novel *looking pattern assay*, designed to assess monkeys’ ability to identify images based on the fixation points of their eyes as they viewed images (Methods). This method of assessing differences in the monkey looking patterns between two images is inspired by attention maps in humans^24^. Briefly, we identified points on the screen where monkeys fixated on images^25^, created attention maps by filling each of these fixation points with Gaussian kernels, and quantified the similarity between eye patterns of any pair of images by correlating the kernel fills between the images (Fig. S3). This correlation in the looking pattern assay yields a single value to assess behavior: **ρ**. If monkeys are shown any two random images, the pattern of eye fixations should only coincidentally overlap, and therefore this assay tends towards a low **ρ** value. However, if a novel and repeat image are compared, this assay should tend towards higher values of **ρ** as monkeys identify and attend to features in the previously seen image from memory^26,27^ and therefore are more likely to scan the same path^28,29^. So when monkeys are shown a novel and then lure image, if the monkey successfully discriminates the lure (a “lure”|lure), we expect the **ρ** to be relatively low compared to when the monkey misclassifies the lure image as a repeat (a “repeat”|lure). In this same sense, when monkeys are shown a novel and then repeat image, if the monkey successfully discriminates the repeat (a “repeat”|repeat), we expect the **ρ** to be relatively high compared to when the monkey fails to identify the repeat (a “lure”|repeat). We can then set a single threshold for **ρ** to classify lures and repeats with the same signal detection theory framework described above (Fig. 1C).

Using the best threshold for separating “lure”|lure images from “repeat”|lure images for each individual session (Methods), the looking pattern assay achieved an average “lure”|lure rate of 0.62±0.02 compared to a “lure”|repeat rate of 0.40±0.02 across sessions (Fig. 1D). In other words, this technique classified 62% of lures as lures (“lure”|lure) and 60% of repeats as repeats based on monkey looking patterns. This yields a d’ of 0.64, a 4.6x improvement over preferential looking, while also achieving this classification level with a higher proportion of trials (62% of lures were identified as “lure”|lures compared to 39% for preferential looking, Fig. 1D). As a control, we tried classifying trials using only the number of fixation points by the monkey on each image, in which we selected “lure”|lure and “lure”|repeat images as those lures and repeats with >1 fixation point (d’=0.4, Fig. 1D). The looking pattern assay remained better at classifying trials (1.6x improved d’), indicating that the looking pattern assay utilizes information from the scanpaths^28,29^ and is not just an artifact of center-bias due to the requirement of the monkey to center fixate before each image presentation (Methods).

Further, to control for the possibility that the differences in **ρ** between novel and lure images do not relate to memory, but the changes or shifts in the salient features between the two images, we quantitatively assessed the differences between each of our novel and lure image pairs using saliency maps^30^ and then plotted these values vs. the **ρ** we found for the monkey between the same pairs (Fig. S4). If our looking pattern assay is driven by differences in saliency (i.e., perception, as opposed to memory as we hope to gauge), we expect smaller values of **ρ** (which indicate more different scanpaths) will correlate with bigger differences between each pair of images. Instead of finding such a negative relationship, there was a slight positive correlation between the change in salience and **ρ** (orange line; correlation = 0.044, p=0.015), as we’d expect if our looking pattern assay measures memory of the novel image and not just perceptual differences between the novel and lure images.

Based on both preferential looking and the looking pattern assay, we show behavioral evidence that monkeys identify both lures and repeats in a large proportion of trials as they perform the pattern separation task. These methods are also able to quantify monkey identification of lures and repeats on single trials, which we utilize in our final analysis to directly compare behavior and neural responses.

### Neurophysiology

We recorded from 216 single units in the CA3/DG region of the hippocampus in the same three monkeys used in our behavioral analysis during 118 sessions (unit breakdown: rhesus BY=91, rhesus I=115, bonnet B=10). Responses across these neurons during image presentation were highly variable (mean firing rate±std for 158 neurons < 10 Hz = 2.2±2.3 Hz; for 58 putative interneurons ≥ 10 Hz.^31^= 23.7±18.0 Hz) with a population sparseness (a^p^=0.37, Methods) similar to previous recordings in monkey CA3 (a^p^=0.33 ^32^). These neurons were recorded throughout the anterior two-thirds of the hippocampus (Fig. S5-6). Estimates of recording locations within each hippocampal subfield were performed by comparing labels in a rhesus macaque brain atlas^33^ to MRI for rhesus BY and bonnet B and a combination of MRI (Fig. S5) and histology (Fig. S6) for rhesus I.

We first asked if our population of hippocampal neurons showed evidence of differentiating lure and repeat trials via firing rates. We took the 110 neurons (69 neurons < 10 Hz = 2.6±2.5 Hz; 41 putative interneurons ≥ 10 Hz. = 25.2±20.1 Hz) recorded for at least 50 lure and 50 repeat trials that the monkey viewed for at least 300 ms each and set up a 10-fold, cross-validated regression classifier^34^ on this pseudopopulation, thereby training each fold on 90% of trials and testing on the held-out 10% (Methods). With this setup, we are essentially looking for hippocampal neurons that generally categorize lure v. repeat images, agnostic to any features of individual images themselves. The classifier was trained separately for each time bin, which accumulated spikes for each of the 110 neurons from a moving window of 200 ms with a 100 ms step size. Note that since monkeys decided how long to look at each image, and only trials with a minimum of 300 ms looking time were used, subsequent time bins after 300 ms decreasingly comprised of image viewing (as per the distribution in Fig. 1C) and increasingly comprised of the inter-trial interval (a blank screen that was shown for at least 1500 ms for all monkeys before fixation to begin the next trial).

Since most neurons in this pseudopopulation were recorded for more than 50 lure and 50 repeat trials (mean trials ± SE: 79.8±1.5 lures and 77.5±1.3 repeats), we randomly subsampled trials from each neuron on every run and determined the prediction accuracy^34^ at each time bin for that subset of trials. Running through this process for 200 permutations allowed us to calculate an average prediction accuracy while also providing empirical error bars at the 5^th^ and 95^th^ percentile prediction accuracy. In order to test for significant classification above chance level, we used a maximum statistic method^35^ that takes into account multiple comparisons across time bins by testing for clusters of prediction accuracy that are unlikely to occur by chance with reference to surrogate data (Methods).

The classifier achieved significant prediction accuracy of lure v. repeat trials from 400-800 ms after image onset (Fig. 2A), indicating that the rate code from the population of hippocampal neurons could differentiate lure and repeat images above chance level. Next, we asked if we could classify novel from lure images as well as novel from repeat images using the same method we used for lure v. repeat images. Indeed, classification of novel v. lure images reached prediction accuracy values similar to lure v. repeat (∼60% at peak), with significant classification from 400-900 ms after image presentation (Fig. 2B). Strikingly, classification of novel v. repeat images reached prediction accuracy close to 80% at peak, with significant classification from 300-1600 ms after image presentation (Fig. 2C).

**Figure 2.**
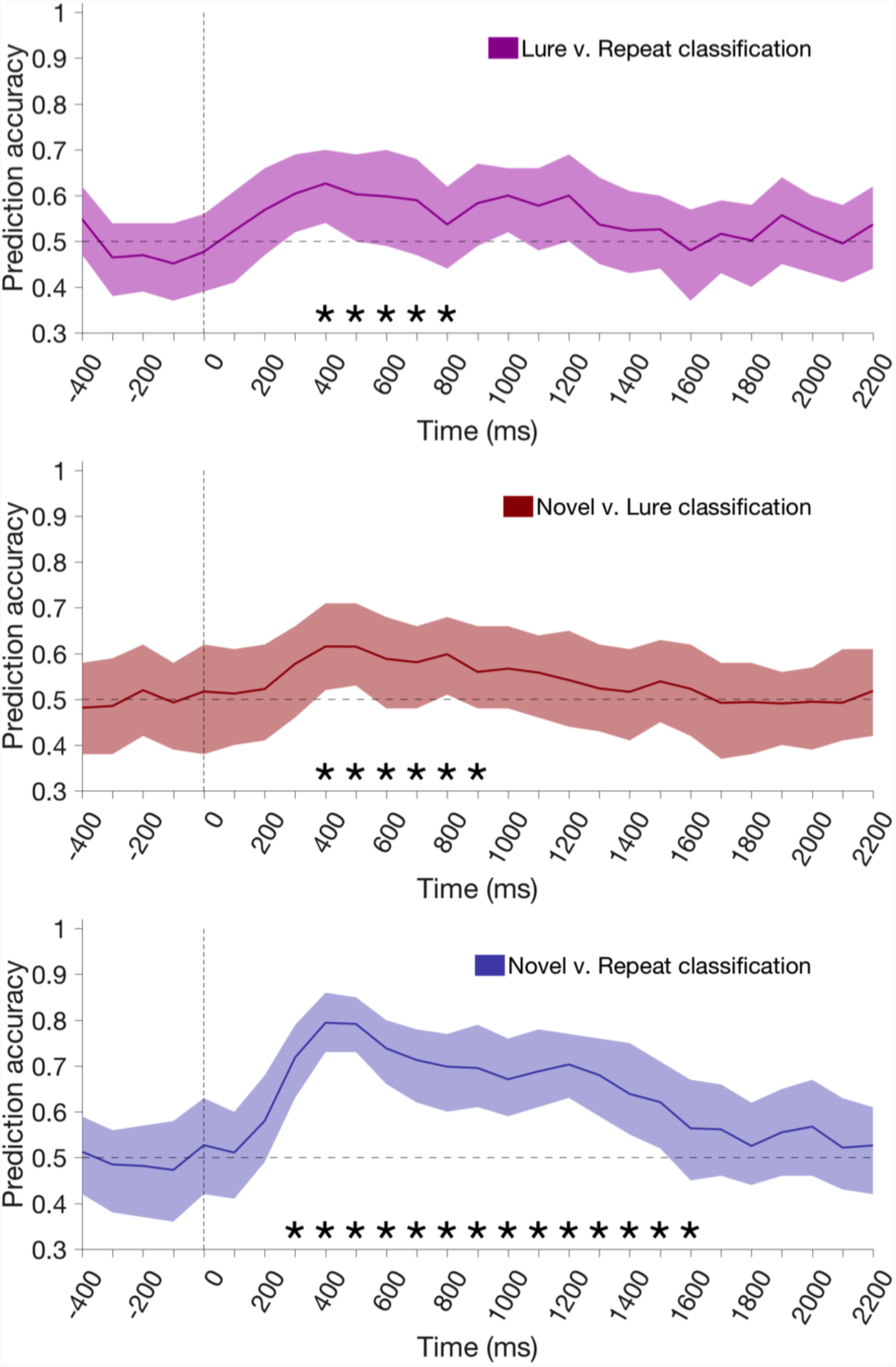
Classification of images by hippocampal neurons. Prediction accuracy from 10-fold cross-validated regression of neural population when comparing **A)** lure v. repeat images **B)** novel v. lure images and **C)** novel v. repeat images. Solid line is average of 40 permutations with different subsampling of trials while errors bars are 5^th^ and 95^th^ percentile prediction accuracy from those 40 permutations. Note that each time point represents spikes accumulated in firing rate windows of 200 ms around that time (e.g. the 300 ms time bin includes spikes from 200-400 ms after stimulus on). Adjacent asterisks in time indicate single, significant clusters of classification above chance level (Methods).

It is surprising that classification of novel v. repeat trials was significant over 1000 ms after image presentation, considering that ∼90% of repeat images were not looked at by the monkey for longer than 1000 ms (Fig. 1B). Could this effect be due to a memory signal being held by neurons beyond stimulus offset and into the (1500 ms) inter-trial interval^36^? To test this possibility, we ran the same classifier but limited images to those with looking times in the range of 300-1000 ms. While the neural population was still able to significantly classify novel from repeat images within 400-800 ms of image presentation, classification returned to chance level (50%) beyond 1000 ms (Fig. S7C). When we carried out the same process for lure v. repeat and novel v. lure images, prediction accuracy still approached 60% within 400 ms of image presentation, but once again returned to chance level beyond 1000 ms (Fig. S7A-B). Therefore, classification many hundreds of ms after image presentation appears to be driven by the subset of trials where the monkey looked at the image for at least that long.

Having shown that we can differentiate the three image types from each other above chance level using a pseudopopulation of hippocampal responses, we investigated the individual neurons that drive these effects. In our classification scheme, the prediction accuracy is achieved by weighting contributions from the 110 neurons in our population for each separate classification shown in Fig. 2. To find the neurons most responsible for all pairwise classifications, we ranked the neurons for each of the three tests by the value of their coefficients when classification was run on a single time bin from 200-700 ms. Neurons with values closest to the top (i.e., the most positive regression weights) for all three tests were selected using a simple penalization method by rank (Methods).

Spike density functions for the top 6 neurons with positive regression weights from this process are shown in Fig. 3A (rasters are in Fig. S8A), while the next 6 are shown in Fig. S9A. Each of these neurons shows a remarkably consistent pattern of responses: firing rates to novel images are the highest, repeat are the lowest, and lure are in between. The firing rates almost always peak within 500 ms of image presentation, and usually return to pre-stimulus firing within 1000 ms (note that the long tail of some neurons is likely due to images that the monkey looked at for that length of time, as explained in the above population analysis and shown by Fig.S7). 3/12 of these neurons were putative interneurons (≥10 Hz baseline firing rates, Wirth, Suzuki), with an average baseline (fixation period) firing rate of 15.9±2.1 Hz, and the remaining 9/12 neurons at baseline of 2.2±1.7 Hz. The response rate above baseline of these 12 neurons was 154±52% during novel trials, 110±39% during lure trials, and 67±33% during repeat trials, each for the 200-700 ms after image presentation.

**Figure 3.**
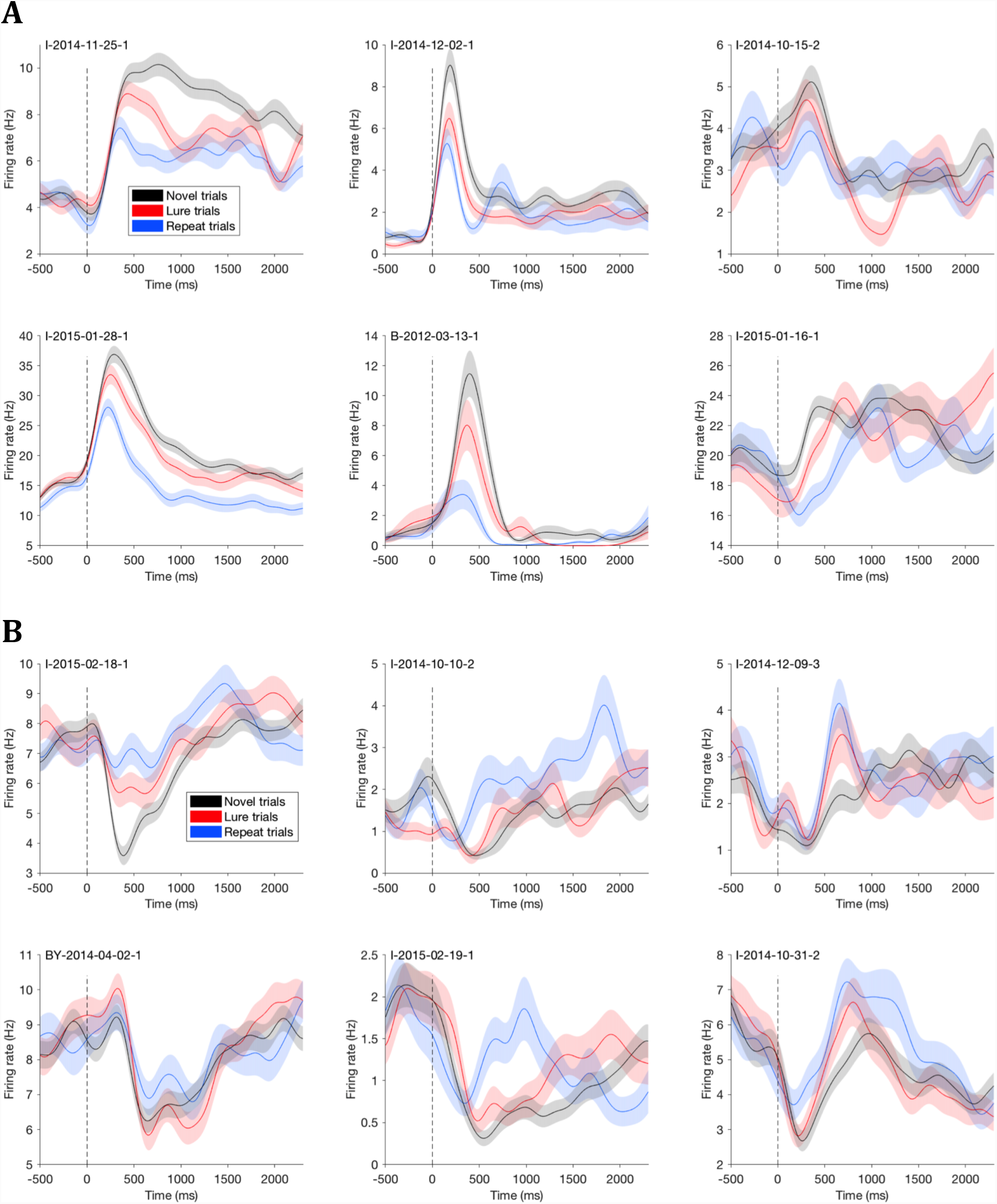
Spike density functions of neurons that contribute most to image classification. **(A)** The six neurons with the highest positive regression weights (in order of weight) from the classifier. **(B)** The six neurons with the lowest negative regression weights (in order of negative weight) from the classifier. Note that both positive and negative weights contribute to the prediction accuracy of the classifier, while neurons with 0 weight do not contribute. The dashed line represents image onset. The spike density functions (solid lines) were calculated by averaging across each trial type after convolving a Gaussian kernel with σ=100 ms for each spike in peri-stimulus time histograms. Shaded regions represent ±1 standard error across each separate trial type. Rasters for every neuron here are shown in sig. S8, while rasters for neurons with the next 6 most positive and next 6 most negative regression weights are shown in Fig. S9. An additional 6 neurons that qualitatively resemble each (positive and negative weight) group are shown in Fig. S10.

We performed the same sorting process for negative regression coefficients for the three pairwise comparisons. These neurons also contribute to prediction accuracy, although the negative coefficients indicate that the pattern of firing rates must be in the opposite direction as neurons with positive weights. The top 6 neurons from this process are shown in Fig. 3B (rasters are in Fig. S8B), while the next 6 are shown in Fig. S9B. Again, this second group of neurons shows a consistent pattern of responses: firing rates to novel images are the lowest, repeat are the highest, and lure are in between. The majority of these neurons also show an initial inhibition of firing within 500 ms of image presentation typically followed by a gradual rise back to pre-stimulus levels hundreds of ms later. Additional examples with qualitative similarities to each (positive and negative regression weight) group are shown in Fig. S10. 3/12 of these neurons were putative interneurons, with an average baseline firing rate of 12.3±2.9 Hz, and the remaining 9/12 neurons at baseline of 2.9±2.2 Hz. The response rate of this second group of 12 neurons was 18±12% below baseline on novel trials, 8±14% above baseline on lure trials, and 13±11% above baseline on repeat trials, each for the 200-700 ms after image presentation. Notably, these neurons with the negative coefficients contributed to the population classifier similarly as the positive coefficient neurons. For example, while the sum of the absolute weights of the 10 most negative neurons was stronger than the sum of the weights for the 10 most positive neurons for all three comparisons, in all three regressions the positive regression weights were within 20% of the absolute sum of the negative regression weights, indicating both groups contribute substantially to the three classifiers.

To quantify the difference in peak firing rate time between the two groups of neurons, we took the top-12 positive regression weight and top-12 negative regression weight neurons and found where firing peaked from 200-1200 ms after image presentation. The distribution of peaks was significantly different between these groups (Kolmogorov-Smirnov (K-S) test, p=9.1E-4), with the positive weight neurons tending to peak a few hundred ms after image presentation, and the negative weight neurons tending to peak closer to 1000 ms after image presentation (Fig. S11).

Is there a difference in how these two groups of neurons classify images? While we formed the groups by finding neurons that had the most positive and negative regression weights 200-700 ms after image presentation for all three of our pairwise classifications, do both groups actually contribute to prediction accuracy for all three comparisons? To test this, we ran the same cross-validated regression as in Fig. 2, but separately for the 12 neurons with the most positive regression weights and the 12 neurons with the most negative regression weights. For the positive regression weight group (Fig. 3A & Fig. S9A), these neurons are still able to significantly classify all three pairwise comparisons within 500 ms of image presentation at ≥60% prediction accuracy level (Fig. 4A). However, for the negative regression weight group (Fig 3B & Fig. S9B), while novel v. repeat (p=0.11, maximum cluster permutation test, Methods) and novel v. lure (p=0.11) classification showed similar levels of significant prediction accuracy as the positive weight group, the classification of lure v. repeat images was significantly greater for the positive regression weight neurons than the negative regression weight neurons (Fig. 4B, p=0.007). Therefore, the classification of lure v. repeat images within a few hundred ms of image presentation seen in our full population classifier (Fig. 2A) is driven by the first group of neurons with positive regression weights.

**Figure 4.**
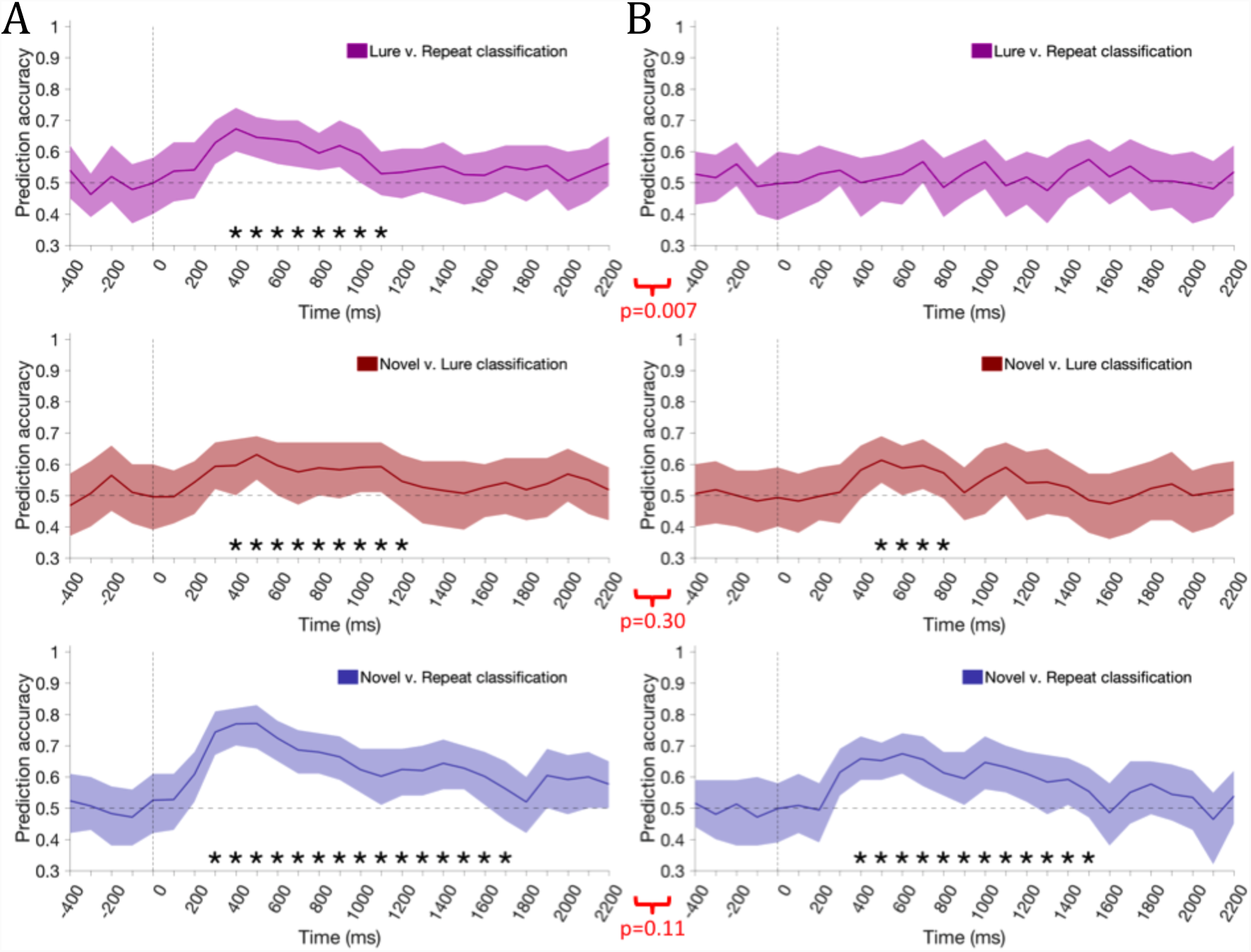
Classification of images by only neurons from positive or negative regression weights groups. Prediction accuracy from the same 10-fold cross-validated regression as in Fig. 2, but using only **(A)** the 12 neurons with the most positive regression weights or **(B)** the 12 neurons with the most negative regression weights. Conventions and statistics for individual graphs as in Fig. 2. Comparisons between graphs used a maximum cluster—based permutation test to calculate an empirical p-value (Methods).

As we’ve shown the ability to classify each of our images from each other with as few as 12 neurons, we asked if we could predict image identity from responses at the level of single neurons. We ran cross-validated regression as in Fig. 2, but for each individual neuron shown in Fig. 3. The results are shown in Fig. S12. While there are clear bouts of significant classification for each of the three comparisons for individual neurons, the prediction accuracy is notably noisy. Many bouts of significant accuracy are interrupted by chance level classification within a few hundred ms of image presentation, while nonsensical time bins—like before image presentation—often show significant prediction accuracy. These classification problems were not evident when training the classifier with as few as 12 neurons as in Fig. 4, indicating that even small groups of neurons are capable of denoising the variability of single neurons.

These two groups of neurons that respond differentially to novel, repeat, and lure images show a stereotyped pattern 1) the 1^st^ group rises in firing rate within 500 ms of image presentation with novel images showing the highest firing rates and repeat images showing the lowest firing rates, and 2) the 2^nd^ group drops in firing rate in this same period before returning to baseline levels with novel images showing the lowest firing rates and repeat images showing the highest firing rates. We asked if this pattern of responses, when pooled across our pseudopopulation, would match the BOLD responses from fMRI found in CA3/DG during pattern separation tasks similar to ours^2,5^. In these studies, repeat images show reduced BOLD responses compared to novel images while lure images show levels of BOLD closer to that of novel images^2,3,5^. In essence, this BOLD response resembles the firing pattern of the 1^st^ group of neurons, as if the higher firing rates of the first group overwhelm the 2^nd^ group when looking across the population.

To simulate a BOLD-like response from the population firing rates of our neurons, we assume the 110 neurons from our classification analysis to be a reasonable sample of the active neurons throughout the hippocampus, and that any given trial in any given session is on equal footing. In this case, we can combine all trials across all neurons into large pools of trials split by trial type (total trials recorded from each trial type: 26396 novel, 9080 lure and 9056 repeat trials). Pooling trials in this manner—after finding the firing rate for each trial from 200-500 ms after image presentation and normalizing it to the fixation period (baseline) firing rate for that neuron—is a simulation of how 26396 hippocampal neurons would simultaneously respond to a *single* novel trial, 9080 neurons to a *single* lure trial or 9056 neurons to a *single* repeat trial.

To visualize how the population firing rates are distributed amongst each group of trials, we created quantile-quantile (QQ) plots^37^ of the firing rate counts for both lure v. novel and repeat v. novel trials (Fig. 5). QQ-plots provide a way to compare how two sets of data are distributed, with a deviation from the line at y=x in a given range representing a difference in the proportion of one distribution over the other within that range. Comparing lure firing rates with novel firing rates, the QQ-plot shows little deviation between the populations as the line hugs y=x until novel firing rates exceed a z-score of 3 (Fig. 5, red). Indeed, the lure and novel QQ distributions do not show a significant difference (p=0.47, K-S test). However, repeat firing rates show a relatively larger difference from novel firing rates, as the proportion of firing rates begins to drop above a z-score of 2 and continues to decline as the firing rates get larger (Fig. 5, blue). This effect is confirmed statistically, as the novel and repeat QQ distributions are significantly different (p=9.3E-4, K-S test), and notably, the lure and repeat QQ distributions also show a significant difference (p=0.021, K-S test). In sum, this analysis shows that the pattern of BOLD responses seen in human fMRI—a decrease in BOLD during repeat images vs. relatively similar levels of activity between lure and novel images, might be explained by the net activity of neural firing rates from image-responsive neurons like those shown in Fig. 3A.

**Figure 5.**
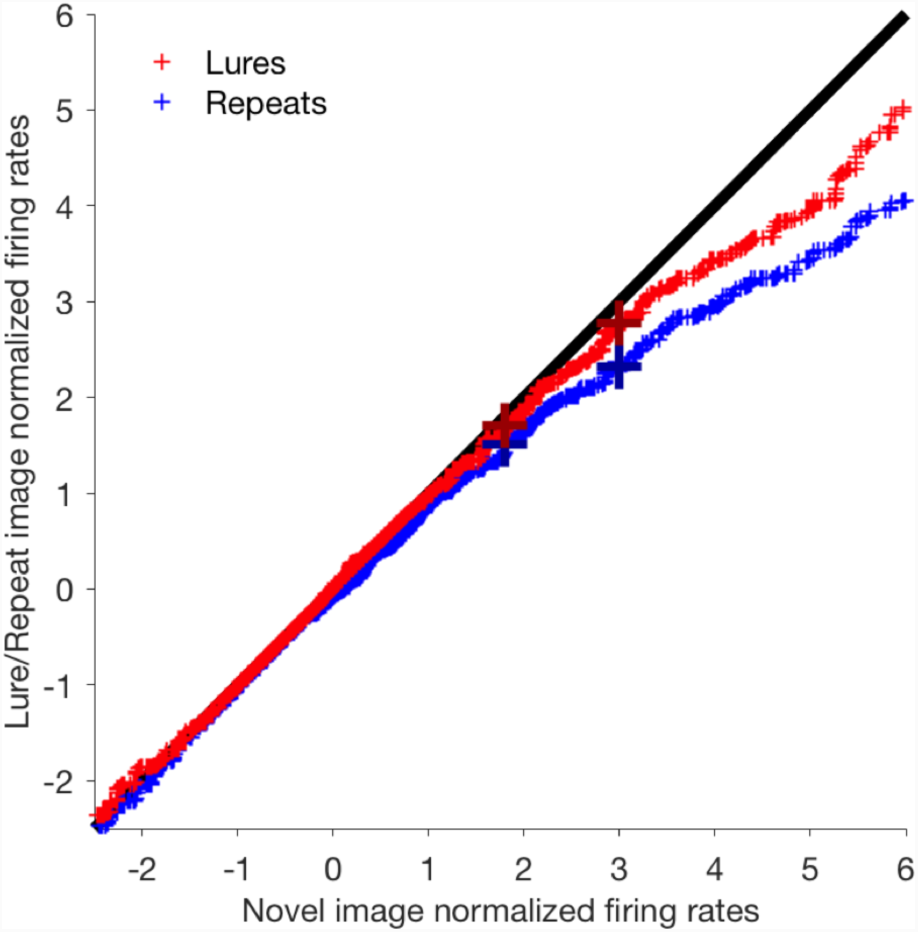
Firing rates across the neural population are lower for repeat than lure and novel images. After pooling the normalized firing rates for all trials for each trial type across all neurons, we created QQ plots of novel v. repeat trials (blue) and novel v. lure trials (red) to visualize the responses to each trial type across the population. The large crosses represent the trial at which 90 (left) and 95% (right) of repeat (dark blue) and lure (maroon) trials are below that normalized firing rate.

To this point, we have shown behavioral evidence that monkeys can identify lure and repeat images, and also classified trial types based on firing rates from groups of hippocampal neurons. Next, we asked if we could use our ability to behaviorally assess image recognition on single trials to confirm the pattern of firing our classifier used to predict trial types. In other words, when we have stronger (behavioral) evidence that the monkey identifies a particular image type, do we see a stronger (neural) response in support of that identification?

The neurons most responsible for image classification showed the most drastic change in firing rate to repeat trials as compared to novel trials: either lower firing to repeats for the 1^st^ group of neurons (Fig. 3A) or higher firing to repeats for the 2^nd^ group of neurons (Fig. 3B). Meanwhile, the change in firing rates between lure and novel images was relatively smaller than the change between repeat and novel images for both groups (i.e., lure firing is almost always between novel and repeat firing for all neurons in Fig. 3 & Figs. S8-10). Therefore, our expectation is that on a lure trial where our behavioral assay indicates the monkey is more likely to have identified the lure image as a lure than a repeat image (lower **ρ**), we expect a relatively smaller change in firing rate between the lure and novel image. On the other hand, when our behavioral assay indicates the monkey is more likely to have mistaken the lure image for a repeat image (higher **ρ**), we expect a relatively larger change in firing rate between the lure and novel image. We can test this expectation for each neuron by correlating the change in firing rate during the presentation of each lure image and its novel image pair (ΔFR) with the output from our behavioral assay (**ρ**) for each pair of images (Figure 6A). On these graphs, positive correlations for lure trials provide evidence of our expectation: that CA3/DG neurons maintain more similar levels of firing between novel and lure images (low ΔFR) during successful pattern separation of lure images (low **ρ**).

**Figure 6.**
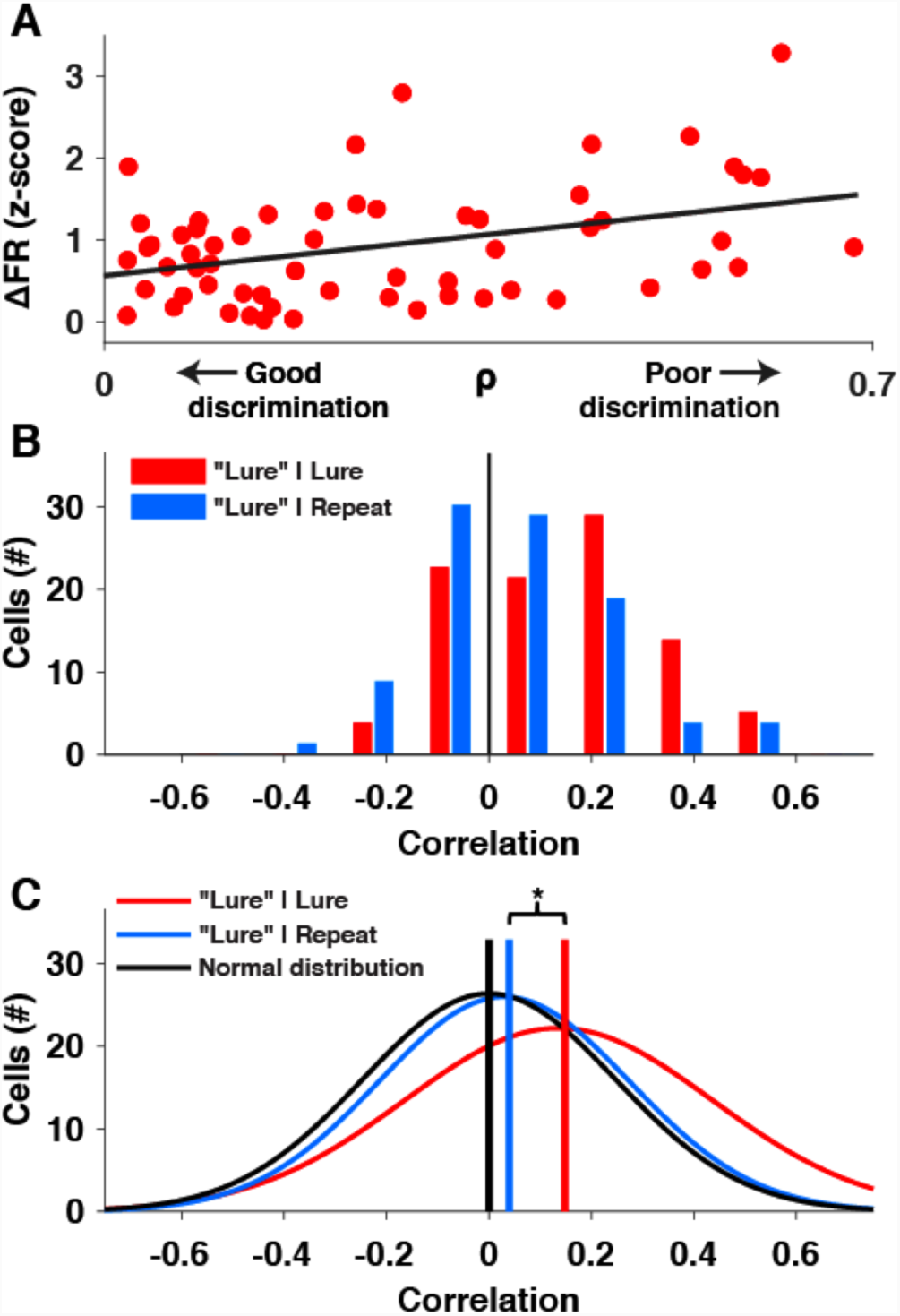
Neural firing rates correspond to behavioral pattern separation for single cells. **A)** Scatterplot for an example neuron, where each point shows the change in firing rate (ΔFR) v. the looking pattern assay value **ρ** for a single novel and “lure”|lure image pair. Black line is a linear fit of the points, which have a correlation of R=0.43 (p=3.3e-4, Pearson). Positive R values indicate more similar firing rates (smaller ΔFR) during better behavioral pattern separation (smaller **ρ**). **B)** Histogram of the correlation values (R) between ΔFR and **ρ** for “lure”|lure (red) and “lure”|repeat (light blue) trials for these 76 neurons. Black line is at x=0 for visual effect. **C)** Gaussian fits of each set of data in **B**, along with a null Gaussian distribution (black) for visual comparison centered at 0 with σ averaged from the 2 datasets. The vertical lines indicate the median of each distribution. *“lure”|lure v. “lure”|repeat, p=0.0043 (multi-level ANOVA, Methods).

We found the correlation, R, between ΔFR and **ρ** (as in Fig. 6A) for “lure”|lure trials for each of the 76 neurons with at least 20 “lure”|lure and 20 “lure”|repeat trials and a minimum baseline FR of 0.5 Hz (more strict selection criteria were used than before due to the sample size needed to form each plot as in Fig. 6A). The histogram of R values for these neurons shows a significantly positive skew—as compared to the null expectation of a population centered at zero—for “lure”|lure trials (Fig. 6B, red; median=0.15, p=1.2E-4, sign test). As a control, we can compare these values to a histogram of R values created using the same method from “lure”|repeat trials (Fig. 6B, blue), which are when monkeys misclassified a repeat as a lure, an indication that the animal did not form a strong memory of the novel image and therefore failed to recognize when it was repeated. In this case, neurons should respond more like they were shown random pairs of novel images, which would make the relationship between ΔFR and **ρ** uncorrelated (R values of 0). Consequently, if the pattern of firing we expect is a signature of behavioral lure identification, we should see higher R values for “lure”|lure trials than “lure”|repeat trials. Indeed, for these same 76 neurons the R values for “lure”|repeat trials were significantly lower than “lure”|lure trials (median=0.044 v. 0.15, Fig. 3B; p=0.0043, multi-level ANOVA (Methods)). These data indicate that the better behavioral evidence we have that a monkey identified a lure image, which we consider a signature of successful pattern separation, the more likely it is that CA3/DG neural firing rates to those lure images approach the level of firing rates to novel images.

## Discussion

We recorded from single neurons in the CA3/dentate gyrus region of the monkey hippocampus as animals serially viewed novel, similar (lure), and repeat images. Monkeys were engaged with this implicit pattern separation task, as evidenced by both looking times (Fig. 1B) and looking patterns (Fig. 1D). Using the population of CA3/DG neurons, we could classify novel, lure, and repeat images from each other using firing rates from these neurons during image presentation (Fig. 2), indicating that hippocampal neurons contain a readout of the monkeys’ discriminability of these images. The neurons responsible for this discrimination could be divided into two groups by their pattern of firing rates (Fig. 3), with only one of these groups capable of distinguishing lure from repeat images (Fig. 4). Finally, by combining behavior with neurophysiology, we find that when monkeys are worse at distinguishing lure images from novel images, the difference in firing rate between these image types increases (Fig. 6), confirming the pattern of firing used in the classification analyses (Figs. 2-4) and predicted by human fMRI studies (Fig. 5). These findings suggest a neural signature of pattern separation within the CA3/DG region of the hippocampus.

It is important to note that when we use the term “pattern separation”, we refer to the process that allows the monkey to discriminate lure images from novel and repeat images. Since we cannot separate CA3 from DG neurons and do not have recordings of areas upstream of CA3/DG—as has recently been done in rodents^10^— we cannot quantify the computation that decorrelates similar inputs (lures) in the spirit of Marr^1^. However, we based this paradigm directly on a task developed to elicit pattern separation behavior in humans^2-5,13^. These studies found a signature BOLD response unique to CA3/DG in contrast to the surrounding medial temporal lobe (i.e. CA1, entorhinal cortex, perirhinal cortex, etc.), where BOLD activity changes most during repeat images, while staying relatively more similar between novel and lure images^2,3,5^. This neural signature was therefore the basis for our initial hypothesis: that CA3/DG neurons should respond to lure images more like novel images than repeat images. Indeed, the first group of neurons with positive regression weights showed this pattern of results, with firing rates strongest to novel images, weakest to repeat images, and lure images in between (Fig. 3A). And since this group of neurons peak in firing rate a few hundred ms after image presentation as the second group (with negative regression weights) are inhibited until they return to baseline firing rate hundreds of ms later, our data is consistent with the pattern of BOLD responses seen in human fMRI studies deriving from the firing rates of the first group of neurons.

Another way we visualized pattern separation at the population level was to combine all novel, lure, and repeat lure trials across all neurons into three single pools of trials (Fig. 5). Since the hippocampus shows trial classification agnostic to the content of any given image, as evidenced by the average pattern of firing rates between novel, lure, and, repeat images for those neurons with the largest absolute weights in our classifier (Fig. 3), we treated these pools as if every trial came from a separate neuron simultaneously responding to a single image. That is, since we consider our population of 110 neurons to be a random sample of the active CA3/DG population, we treat the 9080 lure trials from these neurons as a surrogate for how 9080 neurons would respond simultaneously to a single lure image. This analysis showed the same signature of firing as the first group of neurons (Fig. 3A), with responses to novel images greater than repeat images, and responses to lure images in between. Therefore, this analysis is consistent with neurons from this first group driving the global pattern of responses seen in human fMRI studies^2,3,5^.

Electrophysiological studies of recognition memory in humans^38,39^ and monkeys^21^ have shown evidence of hippocampal neurons that either increase in firing rate to novel images (relative to repeat images) or increase in firing rate to repeat images (relative to novel images). The two groups of neurons we found via our classification analysis replicate these findings, as the first group with positive weights have higher firing rates to novel than repeat images (Fig. 4A), while the second group have more inhibited firing rates to novel than repeat images (Fig. 4B). One previous study compared these neuron types and found both showed evidence of changes in firing correlating with human-reported recognition memory^39^, similar to how we can classify novel v. repeat images using both types of neurons (Fig. 4, bottom). However, to the best of our knowledge, none of these studies have found a difference between what these neurons classify. Our use of lure images in addition to novel and repeat images has elicited a difference: only the first group (novel > repeat firing rate) was able to classify lure from repeat images, while the second group (repeat > novel firing rate) could not (Fig. 4, top). This difference begs the question: what is the function of the second group of neurons? Since this group was still able to discriminate novel images from both lure and repeat images (Fig. 4B, middle and bottom), it’s possible these neurons are part of a novelty recognition circuit upstream or orthogonal to the pattern separation discrimination. Unfortunately, we do not have the anatomical precision with our recording methods to determine the hippocampal subfield (CA3 or DG) or neural subtypes of each group. As this second group is almost exclusively inhibitory during image presentation (Figs. 3B and S9B), it’s possible these neurons represent interneurons within the dentate gyrus^40^. Meanwhile, the first group of neurons, which do show lure v. repeat discriminability, might be CA3 neurons, since this region is thought to be the receiver of pattern separated outputs^1^.

Evidence for a sparse distributed code^32^, in which each memory is encoded by a small proportion of neurons that each contribute to a small number of memories, has been shown during an episodic memory task in the human hippocampus^37^ as well as during spatial exploration in primate hippocampus^32^. It has been suggested that hippocampal neurons also utilize such a code for pattern separation of images, in which newly presented images are discriminated from recently seen images stored in memory via sparse representations by CA3/DG neurons^41^. Two of our findings support the idea of a sparse distributed code for pattern separation. First, when we pool firing rates to novel, repeat, and lure images across all trials (Fig. 5), differences only develop in the tail end of the firing rate distribution, where ∼8% of lure and ∼7% of repeat trials contribute to the key range where separation is apparent (firing rate≥2, Fig. 2B). The small proportion of trials showing this difference between responses to lure and repeat images suggests that only a small proportion of neurons participate in a given pattern separation or recognition event. And despite only ∼8% of lure trials showing these changes in firing rates from the repeat distribution, 71% of neurons populate these ∼8% of trials, indicating that while a large proportion of our neural population participates in pattern separation only a small proportion of neurons do so on any given trial. Second, additional evidence of a sparse distributed code comes from Figure 6, as while most neurons show evidence of pattern separation (theoretically each neuron above a correlation of 0 for “lure”|lure trials in Fig. 6b), there is a large variability in ΔFR for each neuron (e.g., Fig. 4A). This variability suggests that while an individual neuron responds stronger to some pattern separation events than others, it is only involved in a proportion of pattern separation events. Taken together, these findings in which a large number of neurons contribute to pattern separation, but each neuron only contributes during a proportion of trials, is evidence of pattern separation being achieved via a sparse distributed coding scheme. Studies using multicellular recording from dual regions would be essential to directly^10^ show how the underlying sparse distributed code achieves these computations without relying on the pseudopopulations recorded across days here.

A number of studies have shown evidence of population codes of spatial pattern separation in the CA3 and dentate gyrus regions of rodent hippocampus^7-11^. These studies report ensembles of CA3 neurons that show evidence of pattern separation or pattern completion dependent on task demands^7^, while DG displays more abrupt changes in the population coherence with small changes in context^8,10^. While our results confirm that pattern separation manifests as a population response across a mixture^14^ of CA3/DG neurons^8,10^, our findings are difficult to compare with these key rodent pattern separation studies. Other than the usual difficulty in comparing behavior across species, pattern separation tasks used in rodents typically involve discrimination of a small number of physical environments—with recordings focused on place fields—over the course of minutes^8,10^. And in light of recent work indicating the heterogeneity of the rodent hippocampal populations^14^, it becomes even more difficult to come up with a unifying prediction from these studies to compare with our results. In contrast, the use of monkeys allowed us to emulate image pattern separation tasks used in humans, making our results more directly applicable to human fMRI data due to task similarity. Future studies can work to bridge these gaps between research models with new task design in rodents (including parametric, non-spatial tasks that drive hippocampal responses^42^) and more powerful recording techniques in monkeys and humans. In addition, monkeys provide a research model capable of assessing neural responses to other aspects of pattern separation, such as quantifying the pattern separation function via parametrically rotated stimuli^15^ or altering stimulus presentation times to probe temporal pattern separation^1^.

This study identifies how CA3/DG hippocampal neurons respond as monkeys achieve pattern separation of images from those in short term memory. The key neurons that drive discriminability between novel, lure, and repeat images show a pattern of firing rates consistent with both the signature of pattern separation in CA3/DG of human fMRI studies and the population coding of pattern separation via CA3/DG ensembles shown in rodent studies of pattern separation, with one subpopulation appearing key for the discrimination of lure (similar) and repeat images. Further, our data indicate that pattern separation is achieved via a sparse distributed code from across the population of neurons in the hippocampus, with only a small proportion of hippocampal neurons responding on any given pattern separation event despite a large proportion of the population participating in pattern separation overall. With this single neuron signature of pattern separation and evidence for a sparse distributed code across the population in mind, future studies using multicellular recordings—particularly with simultaneous recordings from brain regions that connect to hippocampus—will be essential to understand how pattern separation emerges from the network effects of neural populations.

## Methods

### Subjects

Two female rhesus macaque (rhesus BY=6.7 kg, age 11; rhesus I=5.7 kg, age 6) and one male bonnet macaque (bonnet B=8.9 kg, age 12) monkey were used for experiments. Animals were surgically implanted with titanium headposts using aseptic techniques under general anesthesia and then trained to be comfortable visually interacting with a computer monitor for juice reward while being headposted during a multiple hour recording session. All procedures and treatments were done in accordance with NIH guidelines and were approved by the New York University Animal Welfare Committee.

### Behavior

Animals were positioned in a primate testing chair (Crist Instruments) and trained to fixate on a 19-inch LCD monitor (Dell) positioned 0.45 m. from center of screen to left eye. Eye position on screen was monitored by an infrared camera at 240 Hz. resolution (IScan, Inc.) and linearly interpolated to 200 Hz. resolution so each time bin of eye position was 5 ms. The task was programmed in Presentation (Neurobehavioral Systems).

### Task

In our main task, monkeys were shown series of images centered on a computer screen. Each trial began with the presentation of a centered red dot of radius 1°. Animals received 0.35mL juice reward for maintaining fixation within a 1.3° radius around the center of this dot for 500 ms. Next, animals fixate on a blue centered dot with the same parameters as the last one and when this ‘pre-fixation’ is maintained for 300 ms, a centered image is presented on the screen. Animals were allowed to view the image for up to 10 seconds but the image was removed—and the trial completed—when monkeys looked outside the bounds of the image with a 1.85° surrounding buffer (that is, fixation did not break until the position of the eye was outside of the actual image dimensions plus a 1.85° rectangular frame). After the image was removed, an intertrial interval of 3.0 s (1.5 s for monkey BY) began until the fixation dot came on for the next trial. We kept images at their native resolution unless their longest dimension was >600 pixels (22.2°), in which case the image was scaled down such that the longest dimension was set to this length. Animals were initially trained on a version of the task with only novel and repeat images (identical to the visual preferential looking task [VPLT]^21^) with no lure images. This was done because images used during neurophysiology were never used again for another recording session and we wanted to save most of the lure image set for recording sessions. In a few sessions before recording the animal performed the full task to acclimate to the presence of lure images.

Each image was either 1) a novel image, which was never before shown to the monkey 2) a repeat image, which was the repeat of a novel image shown to the monkey 1-4 serial positions after the initial presentation or 3) a lure image, which was an image similar to a novel image shown 1-4 serial positions after initial presentation. Examples of lure images are shown in Fig. S1. On any given trial— except the first—there was a 3/5 chance of seeing a novel trial and a 1/5 chance of seeing either a lure or a repeat of a previously shown novel image. Therefore, 1/5 of novel trials were foil images, which were only shown once and therefore were not followed by a repeat of the foil or a similar lure version. For all analyses, foil images were not included as part of the novel images, as even though these were indeed first presentations to the monkey, by comparing only novel images that subsequently had similar (lure) or identical (repeat) pairs we hoped to minimize differences in perception to image content (and maximize the focus on memory). Lure and repeat images were shown within the next 1:4 trials after their associated novel image (0-3 intervening images). Lures and repeats both had an equal proportion of these intervening image numbers in each session.

We compiled a set of 6450 lure image pairs, 6450 repeat image pairs and 6450 foils. This large set of pictures allowed us to perform all neural recording sessions with images the monkeys had never before seen, thereby ensuring this task would be focused on short term memory encoding and retrieval. Images were handpicked from a wide variety of sources and are available for research purposes upon request. Images were collected from the following sources: a previous pattern separation study^2^, ^43^(available at http://www.vision.caltech.edu/savarese/3Ddataset.html), Hemera photo objects 25,000 (Hemera, 1999), ^44^(available at http://fei.edu.br/∼cet/facedatabase.html), ^45^(available at http://vision.stanford.edu/lijiali/event_dataset/), ^46^(available at http://www.image-net.org/), screenshots from webpages, and screenshots taken from Youtube.com videos and DVDs using custom software in Matlab. This provided a wide array of image types which included objects, environments, faces, people, animals, etc. (Fig. S1). To ensure there were no unintentional repeats or overly similar lure images across our 19,350 images, we used Visipics V1.3 (www.visipics.com) to compare images and manually remove those with high similarity.

Despite reward being incidental to the behavior in the task, monkeys showed sustained interest in images throughout each session. As shown in Figure S2a, while there was a slight decline in looking time of novel images near the end of the average session, there was no noticeable decline in looking times for lure or repeat images. We also found that monkeys showed no intent to ‘cheat’ the task by immediately saccading outside the bounds of the image to get to the next trial, as animals looked at 96.4% of all images for at least 250 ms, and saccadic reaction times in monkeys rarely exceed 200 ms^47^).

### Behavioral trial classification

We assessed behavior for animals during the 88/118 sessions where at least 50 lures and repeats were shown to the animals (Rhesus BY=37, Rhesus I=43 and Bonnet B = 8). The remaining sessions showed similar average values for our classification metrics but were excluded for quantification of our assays due to their small sample size (and therefore increased variance).

For our first analysis of behavior, we used looking times as a proxy for memory performance. These times were measured as the time from image presentation until the animal looked outside the bounds of the image (including the aforementioned 1.85° spatial buffer around each image). To test for differences between these looking times based on trial types, we used multi-level ANOVA to compare looking times of novel, lure and repeat images. Our predictors included categorical variables for the three monkeys and the three trial types tested, with the dependent variable the looking time on each trial. Our goal is to test for differences between trial types after regressing out the variance of the looking times between monkeys and the interaction of monkeys X trial types. Therefore, the resulting p-value we report indicates only the significance of the trial type factor. We used the anovan function in Matlab to make this comparison and then used the multcompare function to perform pairwise, multiple comparisons—corrected Tukey HSD tests between pairs of trial types.

To address the possibility that looking times for lure images are between looking times for novel and repeat images because monkeys treat lures as a mixture of novel and repeat images, we compared the variances of pooled lure looking times vs. pooled novel and repeat looking times. For these tests, we kept only images with looking times between 0.2-5.0 seconds to ensure outliers did not drive the difference in variance. We also pooled the repeat images with only the novel images that later were followed by lure images, with the idea that using only this set of novel images would make the best comparison to the pooled lure images due to the very similar image content. The null hypothesis is no difference in the variance between lure looking times and combined novel and repeat looking times. This hypothesis was rejected, as lure looking times had significantly lower variance than combined novel and repeat looking times (p=9.3E-7, Levene’s test), indicating that lure images are not merely treated as a mixture of novel and repeat images. Levene’s test was used instead of an F-test since looking times do not follow a normal distribution and this test is less sensitive to normality. Another way to account for the exponential distribution of pooled looking times (Fig. 1B) is to log-transform the data^48^ and perform an F-test. The variance of looking times was still significantly lower for lure images than combined novel and repeat images (p=0.0028). Similar results were found if we used the novel images that later because repeat images instead of the novel images that later became lure images (p=9.2E-5, Levene’s test; p=0.0044, F-test of log-transformed data).

We used a signal detection theory framework to classify behavior on lure and repeat trials, with the true positive condition indicating the monkey looked longer at a lure image than its previously shown accompanying novel image. We refer to such a trial as a “lure”|lure, as if the monkey responded “lure” to a given lure image. The true negative condition is when the animal looked less at a repeat image than its previously shown accompanying novel image, which we refer to as a “repeat”|repeat. Therefore, false negatives are when a lure was viewed for less time than its accompanying novel image, which we refer to as a “repeat”|lure, and false positives are when a repeat is viewed for more time than its accompanying novel trial, which we refer to as a “lure”|repeat.

As shown in the main text, we achieved better classification performance using a looking pattern assay (Fig. 1D), which used the pattern of eye fixations instead of looking times to compare pairs of trials (i.e. novel v. lure and novel v. repeat). This method is inspired by attention map or visual saliency methods used in humans^24,49^. First, for a given comparison (e.g. a novel image and its paired lure image), we found the fixation points separately for each image. Instead of defining fixation by an arbitrary criterion like a minimum amount of time the fixation remained in one location, we used a k-means—based algorithm which was specifically developed for patterns of fixations from monkeys (ClusterFix http://buffalo-lab.physiol.washington.edu/ClusterFix.html)^25^. We took these fixation points on each image and plotted 2-dimensional Gaussian kernels centered at each point with identical weight for each point. The σ for each Gaussian was set to the standard deviation of the distances from the center of the fixation point of each time bin during fixation. An example is shown in Figure S3, where fixation points from the eyetracking data are shown superimposed over a novel and lure image (top). The Gaussian fill of these kernels is shown on the bottom, which create “attention maps.” Finally, the rows of each attention map were appended into two single vectors and the Spearman correlation (**ρ**) between them was calculated. Novel and lure pairs (or novel and repeat pairs) without at least 1 fixation on each image were discarded, as trial classification for these trials was not possible.

We used this correlation value, **ρ**, to classify trials with a signal detection theory approach similar to what was described for looking times above. If we calculated **ρ** for patterns of eye fixations between two completely unrelated images, which would have two random patterns of eye data (except that they both start in the middle of the screen), we would expect a low **ρ** value. On the other hand, when we show two identical images, we expect the patterns of fixations to be more similar, as animals will be more likely to examine the salient features between the two identical images and yield a higher **ρ** value if they recognize the image as a repeat. This effect of similar eyetraces for repeated stimuli has been shown experimentally in humans ^28,29^, while there is a notable history showing that looking patterns are related to hippocampally-dependent memory^26,27^.

Therefore, when the monkey recognizes a repeat image as similar to a previous novel image, a “repeat”|repeat, we expect a relatively high **ρ** value. And when the monkey fails to recognize a repeat image, a “lure”|repeat, we expect a relatively lower **ρ** value. On the other hand, when the monkey treats a lure image as different from a previous novel image, a “lure”|lure, we expect a lower **ρ** value. And when the monkey fails to identify a lure image as different from a novel image but instead treats it more like a repeat, a “repeat”|lure, we expect a higher **ρ** value. By setting a threshold value of **ρ** we can then classify repeats into “repeat”|repeat or “lure”|repeat and lures into “lure”|lure or “repeat”|lure. To find what threshold value optimally classified repeats and lures for each session, we made an ROC curve using “lure”|lure as true positives and “lure”|repeat as false alarms and parametrically tested values of **ρ** to find one that maximized d’. The median ‘best threshold’ calculated for each session was 0.525 and the mean value was 0.56±0.03 (SE). These best thresholds were used for our behavioral analysis in Figure 1D as well as the neural analyses in Figures 4, S6, S8B and S9.

We also classified repeat and lure trials by just using a single threshold for all sessions. While using a constant threshold worsened our behavioral classification (session average d’=0.39 as opposed to 0.64 for best threshold, Fig. S7), we used this metric when we wanted to more selectively classify recognition trials (“repeat”|repeat). The intuition for this is that the best threshold calculations described in the previous paragraph maximize the difference between true positives (“lure”|lure) and false alarms (“lure”|repeat). This method left us with 40% of repeat trials selected as “lure”|repeat (Fig. 1D). Therefore, the other 60% of repeat images were classified as “repeat”|repeat. To more discriminatingly select “repeat”|repeat trials, when we set a relatively higher single threshold of 0.6 for all sessions, the average number of repeat trials classified as “repeat”|repeat was reduced to 41% (a refinement of 19%). Meanwhile, setting this constant 0.6 value only came at a cost of classifying lures as “lure”|lure at 70% (8% less refined than best threshold, Fig. 1D). We used a single threshold for Figure 3, since we wanted to compare pattern separation (“lure”|lure) v. recognition (“repeat”|repeat) with this more refined selection of repeat trials. However, we still show similar trends when using the less-selective-for-recognition best threshold method (Fig. S8B).

As a control, we created a classification scheme for lure and repeat trials in which we identified the number of fixation points (using ClusterFix, as explained above) for each lure and repeat image and classified lures with >1 fixation point as “lure”|lure and repeats with >1 fixation point as “repeat”|repeat. The rationale for this control is that monkeys are required to begin fixation at the center of each image, so the simplest comparison we can make is between a pair of novel and lure (or novel and repeat) images that each have 1 fixation point in the center before the monkey saccades outside the bounds of each. In other words, an argument could be made that our looking pattern assay only achieves positive d’ by classifying such trials as “repeat”|repeat or “repeat”|lure while the remaining trials with more rich eyetraces are classified as “lure”|lure or “lure”|repeat. We show that while such a system does show a positive d’ value of 0.4, our looking pattern assay improves on this value by 60% to 0.64.

As another control of our behavioral metric, we assessed if differences in salience between the novel and lure images were predictive of our looking pattern assay value, **ρ**. We used the SaliencyToolbox^30^ to create a saliency map—which is computed via a model that accounts for color, luminance and orientation contrast to determine the most salient features in a given image—for each equally sized novel and lure image in our set. These maps create grids from each image (e.g. a 400×400 pixel image is turned into a 25×25 grid) and output the predicted saliency (expected foci of attention) with a single value for each square in the grid. If we create a saliency map from a novel image, and sum its absolute differences from a saliency map of its paired lure image, we can essentially quantify the expected difference in image features weighted between color, luminance and orientation between novel and lure images. More simply, this gives us a method to predict how differently any given novel and lure image are expected to be perceived. Then, for each novel and lure image pair, we can plot the predicted saliency difference (that is, the sum of the absolute differences for each square in the grid) vs. the **ρ** value between the same novel and lure image pair from our looking pattern assay. If our looking pattern assay is driven by differences in saliency (that is, perception, as opposed to memory), we expect smaller values of **ρ** (which indicate more different scanpaths) will correlate with bigger differences in salience.

### Neurophysiology

Circular HDPE plastic recording chambers (Rogue Research) were custom-fit to the skull of each monkey via structural MRI scans, placed perpendicularly to the horizontal plane in stereotaxic coordinates and dorsally above the anterior 2/3 of the left hippocampus. Single unit recordings were performed by lowering glass-insulated electrodes (∼1 MΩ impedance, Alpha Omega) or glass-insulated tetrodes (∼0.5 MΩ impedance for all 4 shafts, Thomas Recording, Germany) via an electric (NAN Instruments, Israel) or hydraulic microdrive (Kopf Instruments) through 23 gauge metal guide tubes that reached from the chamber grid to ∼10-15 mm. dorsal to the hippocampus. Guide tubes were placed at new locations at the beginning of each week and removed at the end of the week after multiple days of recording. Neural signals were acquired, filtered, amplified 1000x and digitized via the MAP Data Acquisition System (Plexon) at 40000 kHz resolution and sorted manually using a combination of offline sorter (Plexon) and superparamagnetic clustering (Wave_Clus http://www2.le.ac.uk/centres/csn/software)^50^.

No attempt was made to pre-select neurons for any criterion other than the first active cells that were encountered as the electrodes reached expected depths of the dorsal extent of the hippocampus on a given day. Behavioral recordings started after single units were isolated and could be held stably for at least 20 minutes. The 216 neurons in our initial analysis characterizing responses rates represent all units that were recorded during presentation of at least 10 lure and 10 repeat trials with a minimum firing rate of 0.25 Hz. We used more stringent criteria for neuron selection for our population analyses shown in Figs. 2, 5, & 6, as these analyses are dependent on the sample size of the number of trials recorded for each neuron. For Figures 2&5, we used the 110 neurons recorded for at least 50 lure and 50 repeat trials of minimum 300 ms and baseline firing rates ≥0.25 Hz. Unit breakdown was rhesus BY=11, rhesus I=115, bonnet B=4, with mean firing rates for 69 neurons < 10 Hz. = 2.6±2.4 Hz. and for 41 putative interneurons ≥ 10 Hz.^31^= 25.1±20.1 Hz.). Note that, despite the low number of neurons for two of the monkeys (BY and B), neurons from each of these monkeys were members of the key 12 positive regression weight (see Fig. 3A, bottom-center) and 12 negative regression weight (see Fig. 3B, bottom-left and Fig. S9B, top-right) neurons in addition to the neurons shown in Fig. S10. For Figure 6, we used the 76 neurons with at least 20 “lure”|lure and 20 “lure”|repeat trials and a minimum firing rate of 0.5 Hz. In addition, for Figure 6 only, to calculate firing rates during image presentation we measured spikes at an offset of 0.08 s after image presentation (and image removal) to compensate for the latency of visual stimuli to reach hippocampus^51,52^. This was not necessary for the remaining analyses, as in Figures 2-4 we plot the graphs vs. real time and in Figure 5 we only accumulate spikes from 200-500 ms after image presentation as done in previous analyses of this type^37^. Note that for both population analyses in Fig. 2 and Fig. 6, we empirically tested combinations of criteria close to each of these values (for minimum trials, minimum firing rate and offset) via grid search to confirm that such small changes in these criteria did not appreciably change the statistics or our conclusions.

Previous work has used a measure of population sparseness to indicate how many neurons in a population respond to a given stimulus^41^. Since our task was specifically designed to avoid conflation of short-term and long-term memory by never showing the same images within or across sessions (the only images each monkey was shown more than once were novel images that became repeats, meaning they saw these images a total of twice), we cannot truly compare how our population of 137 neurons respond to the same stimulus. However, when we average the firing rate during all image periods for all neurons in our task and calculate population sparseness via the equation:

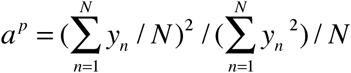

where y_n_ is the average firing rate for each neuron and N is the total number of neurons, we get a value (a^p^=0.37) similar to the population sparseness previously found in primate hippocampus (a^p^=0.33,^32^). When we measured population sparseness this way for more specific image types, values were consistently between 0.36-0.37 (novel, a^p^=0.37; lures, a^p^=0.37; repeats, a^p^=0.37; “lure”|lures, a^p^=0.36; “repeat”|repeats, a^p^=0.37; etc.). Therefore, our data indicate that population sparseness in this range is a general property of the firing distribution of CA3/DG neurons during image presentation regardless of the images shown.

### Population analyses

The first classifier analysis (Fig. 2A) includes the 110 neurons recorded for at least 50 lure and 50 repeat trials of minimum 300 ms and baseline firing rates ≥0.25 Hz. We set up a 10-fold, cross-validated L1-normalized linear regression on these 110 neurons. The cross-validation was balanced, meaning we trained our classifier on 90% of data and tested on the remaining 10% of data for 10 separate iterations, each of which using a different 10% of trials held out on every iteration. We trained and tested a different classifier at each time bin, where each bin accumulated spikes aligned to image presentation for each of the 110 neurons from a moving window of 200 ms with a 100 ms step size. L1-regularized (ridge) regression^53^ was used as with relatively few test trials on each run (10 trials, since 50 lures+50 repeats = 100 trials), perfect classification on some runs led to weights for neurons going to infinity (overfitting). L1-regularization inhibits any given neuron from weighing too heavily in the regression, thereby preventing overfitting while also allowing multiple neurons to contribute to the classifier even if they have redundant signals (we used L1-regularization instead of L2-regularization as the latter attempts to remove such redundant neurons, although classification results were similar for both). For our purposes, we were interested in finding neurons with redundant signals since we wanted to explore what neurons best classify images (Figs. 3-4). The regularization parameter (λ) was set to 0.3, although λ values from 0.1 to 0.5 gave similar classification results.

Since most neurons were recorded for more than 50 lure and 50 repeat trials (mean trials ± SE: 79.8±1.5 lures and 77.5±1.3 repeats), we randomly subsampled 50 of each trial type from each neuron and ran the classifier with these random samples of trials for 200 separate permutations. By running the classifier 200 times with a different subset of trials populating each run, we could then find the average prediction accuracy by averaging across runs. In addition, by rank-ordering the prediction accuracy at each time bin, we could find the 5^th^ and 95^th^ percentile classification runs to create error bars. However, we cannot use these empirical error bars to assess if the prediction accuracy is above chance level at each time bin, as we have to correct for multiple comparisons across time bins. Plus, typically to assess significance for a given comparison, we have to mix across two conditions (e.g. lure v. repeat trials) to create a surrogate distribution, while the error bars above were created through subsampling with the same conditions.

Therefore, to assess significance between conditions while accounting for multiple comparisons over time bins, we used a cluster-based permutation test^35^. This test used the same 200 subsamples of trials as in the last paragraph, but for each subsample the classifier was run after randomizing the labels between the 100 trials (50 trials for each condition). 5 different randomizations were performed for each subsample, creating 1000 total surrogate prediction accuracies at each time bin. For each of these arrays of prediction accuracy over time, a cluster-level statistic^35^ was measured, which is the sum of connected sets of values of those prediction accuracies above a threshold. This threshold can be set to any value without affecting the false alarm rate, but will affect the sensitivity of the test if values are too small or large. We used 0.575, which was just below the average prediction error for all 3 of our comparisons. The sum of the maximum cluster for each of the 1000 permutations was then ranked to create a surrogate distribution. Then, for the prediction accuracy of the real data, clusters are calculated in the same manner, and the rank of the cluster within the surrogate distribution gives an empirical p-value. Since we did three different pairwise comparisons, we consider any cluster less than p=0.05/3=0.0167 (Bonferroni correction) to indicate classification accuracy significantly above chance.

The remaining two classifications (Fig. 2B-C) were identical to the first one, but with novel v. lure and novel v. repeat as the comparisons. The same processes outlined in the last two paragraphs were done for each. Both the average prediction accuracies and significant clusters for all three pairwise comparisons were remarkably consistent across different runs using 200 subsamples of trials, confirming the reliability of this population classifier method despite the random subsampling of trials.

We next determined what neurons were most responsible for each of our three classifications: lure v. repeat, novel v. lure, and novel v. repeat. To do this, we first ran three separate regressions as in Fig. 2, but this time using only a single time bin from 200-700 ms after image presentation. We used this time bin as all three of the comparisons in Fig. 2 show significance in this range. Next, we rank-ordered the vector of regression weights for each neuron in each of the three pairwise comparisons. Finally, we found those cells that contributed most to our three regressions by summing the ranks for all three vectors by cell identity, thereby making the neurons with the lowest sums those that contributed most to the regression via positive weights. For example, neuron I-2014-11-25-1, top left in Fig. 3A, had the 4^th^ highest weight in the lure v. repeat classifier, the 11^th^ highest weight in the novel v. lure classifier, and the highest weight in the novel v. repeat classifier, which sum to 16. The next neuron, I-2014-12-02-1, had the 13^th^, 2^nd^ and 5^th^ ranks, summing to 20, etc.

We performed the same process for the negative regression weights, which also contribute to the classifier, but with the ranks starting from the most negative value. For example, neuron I-2015-02-18-1 had the 6^th^ lowest for lure v. repeat, the 2^nd^ lowest for novel v. lure, and the 3^rd^ lowest for novel v. repeat, summing to 11. The next neuron, I-2014-10-10-2 had the 11^th^, 4^th^, and 2^nd^ ranks, summing to 17, etc.

To compare the contribution of both classes of neurons to each of the three classifiers, we summed the absolute values of the top-10 positive weight and bottom-10 negative weight neurons separately for each classifier. After setting the most positive weight in each of the three vectors to 1 and normalizing all other weights to that value, the absolute sums for the top-10 positive v. bottom-10 negative weights were: 6.8 v. 7.3 for lure v. repeats, 6.1 v. 7.5 for novel v. lure, and 6.1 v. 6.8.

Next, we wanted to see if these two groups of neurons showed differences in classification of images. First, we took the 12 neurons with the highest weights from the positive regression weight group and the 12 neurons with the highest absolute weights from the negative regression weight group and ran the same classifier as we did for the whole population (Fig. 4). Significant classification above chance was tested using the same maximum-cluster method outlined above to account for multiple comparisons over time. However, to assess if one group of neurons performs better than the other at a particular image classification, we required a method that directly compares the classification of each group. To do this, we created a permutation test method using the cluster-level statistic outlined above.

For each of 1000 permutations, we took a random subsample of 100 trials (50 for each condition, as above) for each group of 12 neurons, ran the classifier for those trials, and then calculated the maximum cluster value for each group as described above. Note that we did not shuffle across conditions, as we want to compare the actual prediction accuracies for each group of neurons. Next, we compared the value of these two maximum clusters on every permutation and counted how often one group had a higher value than the other. This creates an empirical p-value for the null hypothesis of no difference in classification between each group, with a p-value <0.0167 (once again, we have to Bonferroni-correct for performing three pairwise comparisons) rejecting the hypothesis that the two groups have similar prediction accuracy for two given images types.

The population spike count analysis (Fig. 5) was modeled after a recent publication on the responses of human hippocampal neurons to newly formed episodic memories^37^. First, we took only images that monkeys viewed for at least 300 ms to match the same trials from the analysis in Figure 2, and measured the spike counts in the window from 200-500 ms after image presentation. This represented 61.4% of all images presented to the animals (Fig. 1B). We then took each image presentation period for each neuron and counted the spikes within this time period and divided by 300 ms to convert to Hz. We normalized the firing rates for each neuron separately by pooling the fixation period firing rates during each session and z-scoring based on the mean and standard deviation of these (baseline) fixation firing rates for each neuron. This allowed us to pool z-scored firing rates across all neurons and then split them into separate groups for each trial type we wanted to compare: novel images (26396 trials), lure images (9080 trials) and repeat images (9056 trials). Quantile-quantile (Q-Q) plots were made using the Matlab qqplot function, which accounts for the different number of trials in both pools to give an equitable comparison of the firing distributions.

Spike counts (and therefore firing rates) for each trial in Figure 5 do not have normal distributions (this was also confirmed by Kolmogorov-Smirnov Goodness-of-Fit test for normal distributions, p<1E-100; kstest in Matlab). In addition, the distributions for each of our firing rate counts had unequal variances (Bartlett test, p<1E-100; vartestn in Matlab). Therefore, we transformed the data to improve the normality and variances of the distributions for each trial type. We found a log transformation does a better job equalizing the variances than the untransformed data or square-root transformed data, as shown by the highest p-values for the Bartlett test^48^. For the log transformed and square-root transformed data, we added a constant value of 5 to each value in the original data to avoid negative numbers, since the minimum value in all 3 distributions was -4.9.

Once we transformed the distributions of firing rates for each of the three trial types, we performed a multi-level ANOVA on the transformed data to test for evidence of differences in the group means between the three distributions. The ANOVA—as well as the following ANOVAs for firing rates tested in this manuscript—is multi-level as we created categorical variables that labeled which of the three monkeys each spike count came from (“monkeys” variable) as well as which trial type each spike count came from (“trial_types” variable). These two categorical variables were then used to predict the vector of firing rates for all trials. This analysis creates three potential factors that could explain the variance in firing rates: differences between monkeys, differences between trial_types and differences between the interaction of monkeys X trial_types. We are interested in testing for differences between trial types after regressing out the variance from the other two factors. Therefore, p-values reported for each multi-level ANOVA test are for the trial type factor after inputting monkeys and trial_types as the categorical variables and the firing rates as the dependent variable using the Matlab function anovan.

Once we determined that there is evidence of differences between the group means between trial types, we tested for pairwise differences in the distributions using a two-sample Kolmogorov-Smirnov (K-S) test. Since we see evidence of differences between the novel v. lure and novel v. repeat distributions when visualized via QQ-plots (Fig. 5), this test is ideal since it quantifies the similarity of two distributions using the absolute maximum distance from the x=y line in the QQ-plot.

For this analysis, we took each neuron and found ΔFR between each “lure”|lure and its paired novel image by aggregating spike counts during the presentation of each image, dividing by the time the monkey looked at that image, z-scoring using the fixation period firing rate for that neuron, and finding the absolute difference in rate between the pair (Δ*FR* = |*z*(*novel*_*FR*)–*z*(*lure_FR*) |, Methods). Since the strength of the correlations between neural firing rates and behavior depend on the number of points populating each scatterplot, we limited this analysis to the 76 single units that were recorded for at least 20 “lure”|lure trials, 20 “lure”|repeat trials and had an average firing rate of at least 0.5 Hz. during all image presentations.

For the analysis in Figure 6 we used only the 76 neurons recorded during at least 20 “lure”|lure trials, 20 “repeat”|repeat trials and with a minimum firing rate of 0.5 Hz. For each neuron, we found the absolute change in firing rate (ΔFR) between each pair of novel and “lure”|lure images as well as for each pair of novel and “lure”|repeat images after z-scoring using the fixation period firing rates for the mean and standard deviation. Spikes were accumulated across the entire time the image was shown on screen and divided by this looking time to convert to Hz before z-scoring. We then found the **ρ** value from our looking pattern assay for each of these pairs of images (novel v. “lure”|lure and novel v. “lure”|repeat) and made separate scatterplots between ΔFR v. **ρ** for “lure”|lures and “lure”|repeats for each of our 76 neurons (example in Figure 6A). The Pearson correlation, which we call R, for each of these scatterplots is shown in Figure 6B. The median value of each is shown with a dashed line. Sign tests were done using the Matlab function signtest with the null hypothesis that the correlations come from a distribution with a median of 0. Comparisons between distributions were done using multi-level ANOVA, where the predictors were categorical variables labeling the monkey that each neuron was from as well as the trial types being compared, while the dependent variable was the correlation R of each scatterplot. In this case, we regressed out the variance from the monkey and monkey X trial_type variables such that we could specifically probe the effect of trial_type on R and therefore report the p-value of this test.

We fit the “lure”|lures, “lure”|repeat and “repeat”|repeat histograms in Fig. 6B with unimodal Gaussians using the Matlab function fit. The black line is a null distribution for visual effect with the average standard deviation of the other 3 histograms and centered at 0.

All data analyses were done in Matlab using custom scripts.

## Data availability

The data and analysis code in support of the findings are available from the lead author on reasonable request.

## Acknowledgements

We thank Craig Stark for helpful comments on a previous draft of the manuscript, John Wixted, Yuji Naya, Shih-pi Ku, and the Kiani Lab for helpful discussions, Arielle Tambini for pilot work on an early version of the project, Andrea Shang and Ellen Wang for assistance in animal training, Shohan Hasan for assistance in image collection and task programming, and anonymous reviewers for suggestions to improve statistics. This work was supported by NIH grant MH084964.

## Author contributions

J.J.S. performed research and analyzed data. J.J.S. and W.A.S. designed research and wrote the paper.

## Competing financial interests

The authors declare no competing interests.

**Population coding of pattern separation in the monkey hippocampus**

## Supplementary Figures (S1-S11)

**Figure S1:**
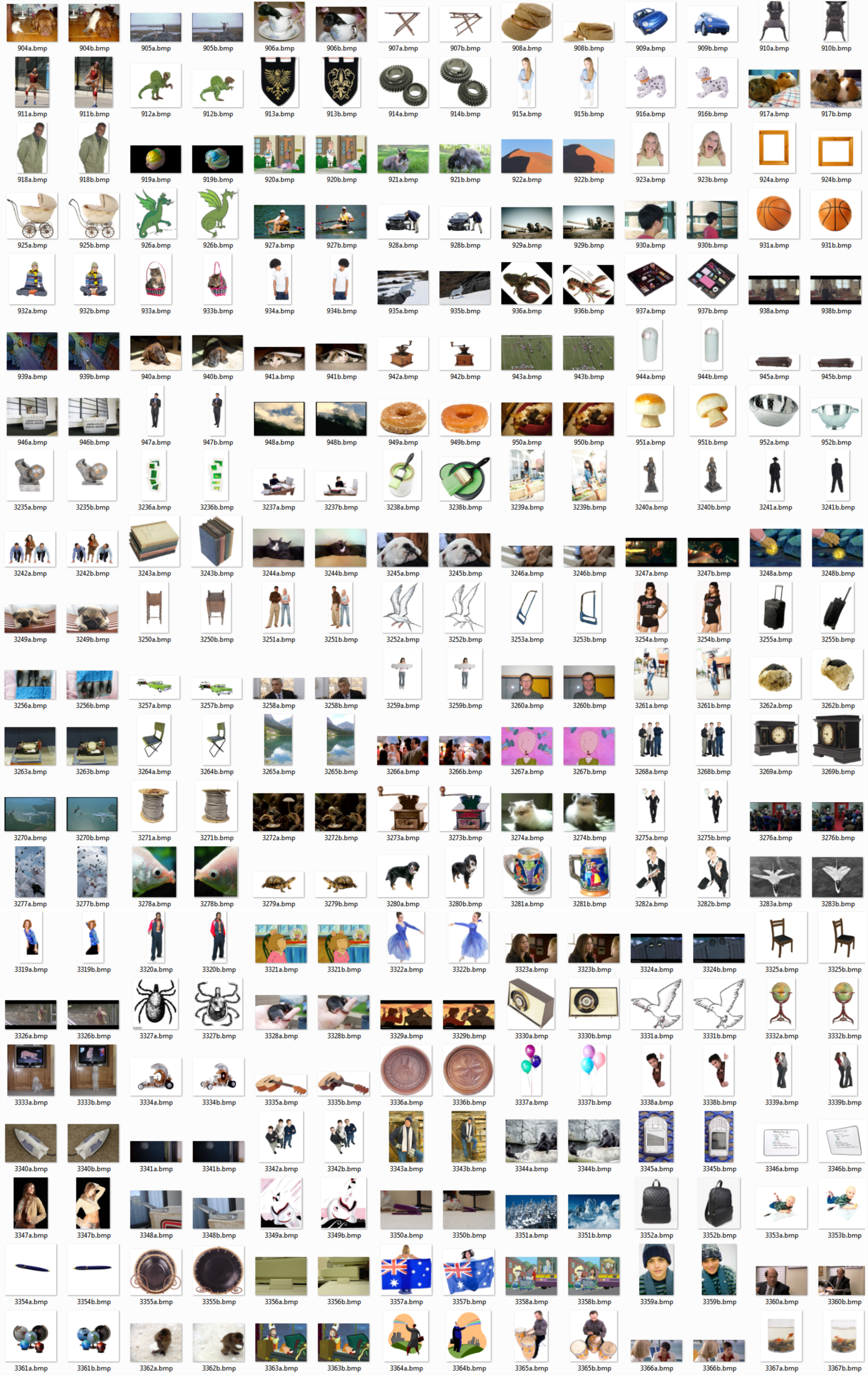
Example images. Randomly selected example novel (“a”)and lure (“b”) image pairs.

**Figure S2:**
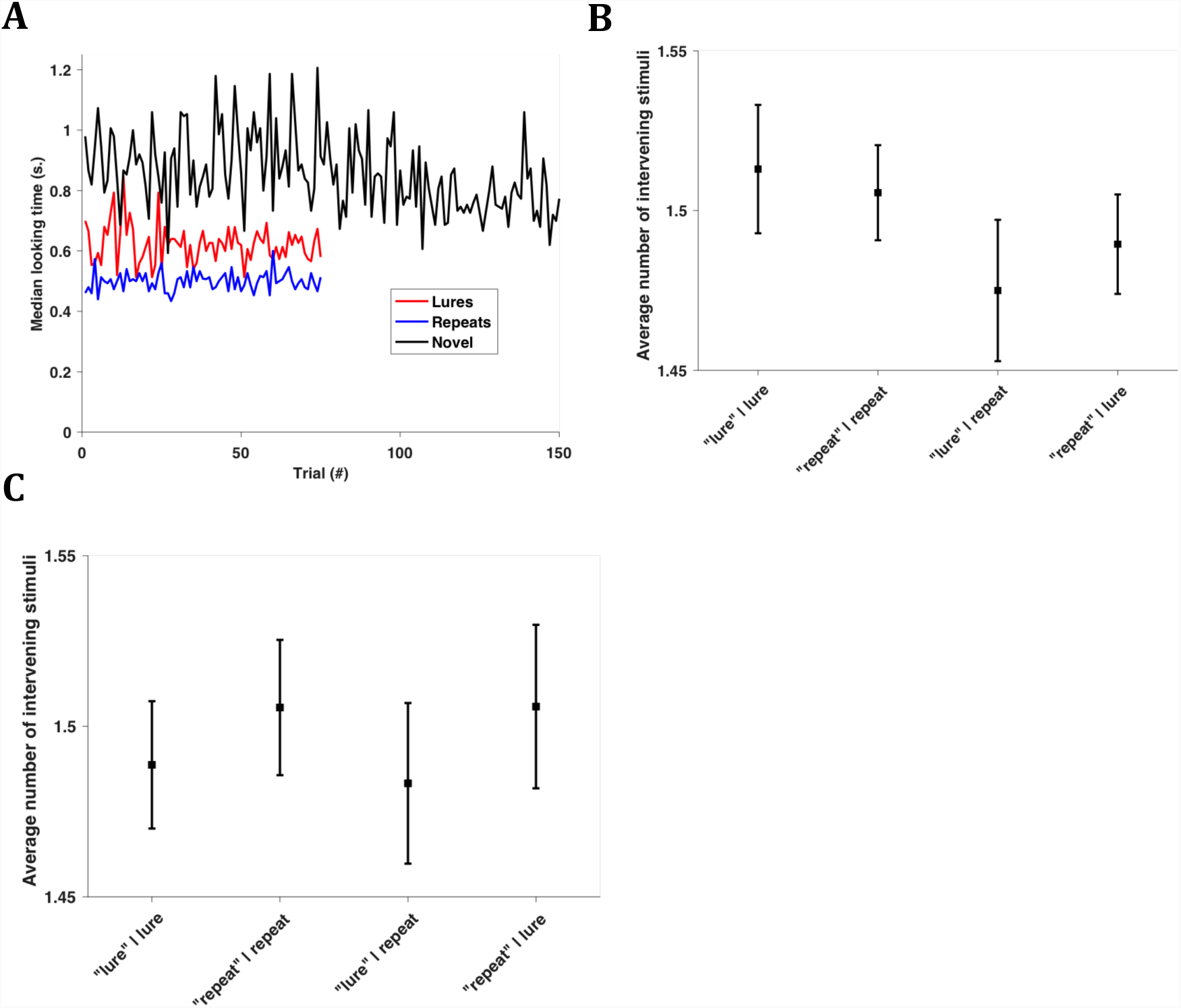
Tests of behavioral performance. **A)** For the 62 recording sessions with at least 150 novel, 75 repeat and 75 lure images, we found the median looking time at each trial position to see if image looking times decrease during sessions (mean values had similar trends but are noisier due to outlier times). As shown, there is no decrease in looking times for lure (red) or repeat (blue) images, which maintain looking times around 600 and 500 ms, respectively. Novel image looking times decline after about 100 novel images were shown. Note that the 100^th^ novel image would be the 250^th^ trial shown in the average session (2/5 of total images shown are novel). The 250^th^ trial in the average session would be where the 50^th^ novel and lure images are shown (1/5 images are lures and 1/5 are repeats, with 1/5 foil images the remaining). Notably, even though there is a decline in looking times for novel images after the 100^th^ novel image is shown, there is no concomitant decrease in looking times for lure or repeat trials after the 50^th^ trial is shown. **B)** We pooled all trials for each classified trial type across sessions and asked if monkeys were better at recognizing lure and repeat images when there were less intervening stimuli between the novel and lure or novel and repeat image. The range of intervening stimuli between each lure or repeat pair was between 0 and 3. Our expectation would be that “lure”|lures and “repeat”|repeats (that is, successfully identified lures and repeats) should on average have less intervening stimuli. This was not the case, as the average number of intervening stimuli for each trial type was not statistically different whether using looking times (p=0.82, multi-level ANOVA) or **C)** the looking pattern assay (p=0.63, multi-level ANOVA) to classify trials. Each of the multi-level ANOVAs used categorical variables for the three monkeys and the four classified trial types as predictors with the number of intervening stimuli for each trial as the dependent variable. P-values represent the trial type factors after regressing out the variance between monkeys and the interaction between monkeys and trial types.

**Figure S3:**
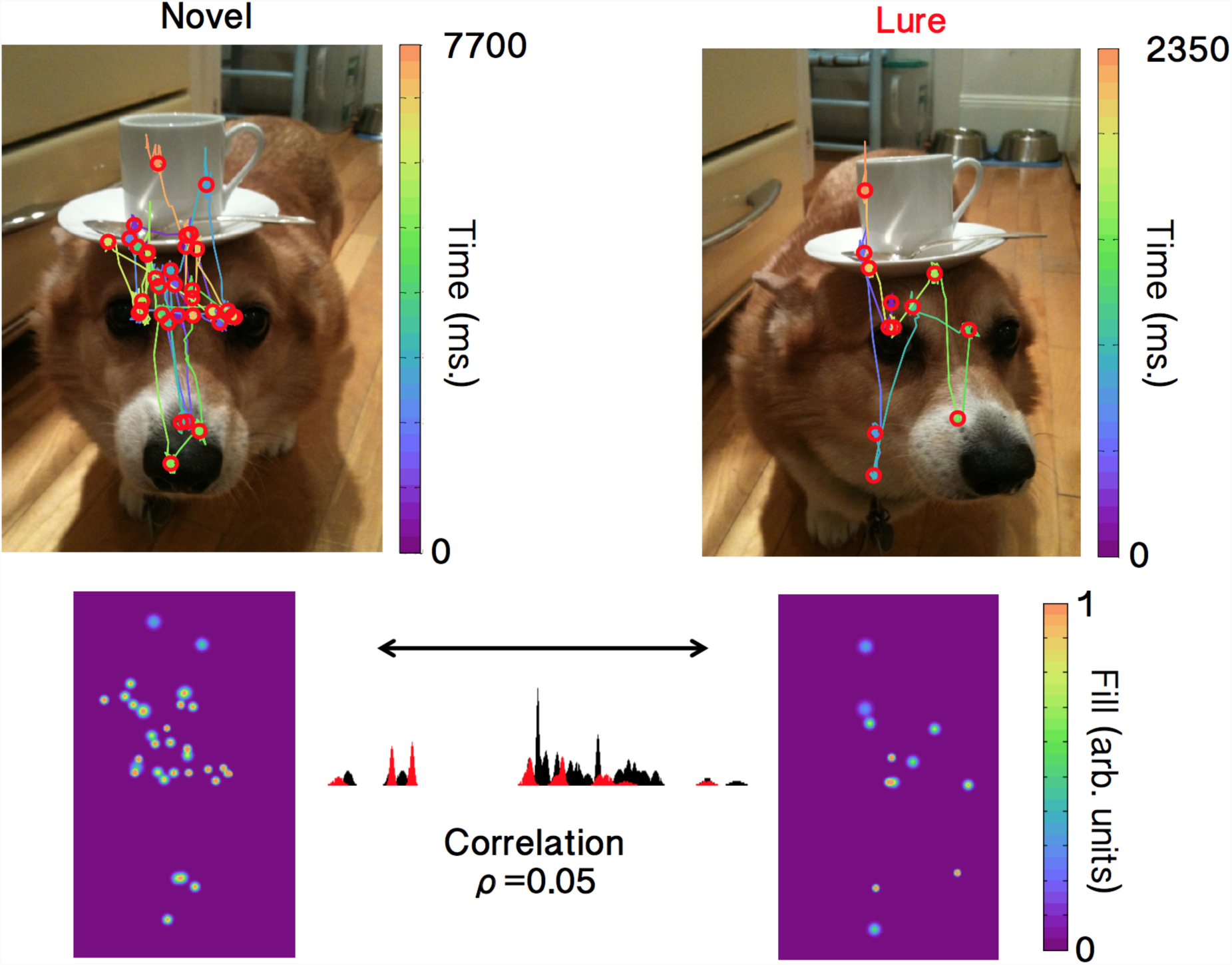
Example images and Gaussian kernel fills for looking pattern analysis. Fixation points were found using ClusterFix^25^. Gaussian kernels were added at each detected fixation point and the concatenated rows of the heat maps (bottom images) were correlated to get a single measure of similarity, **ρ**, between the images.

**Figure S4:**
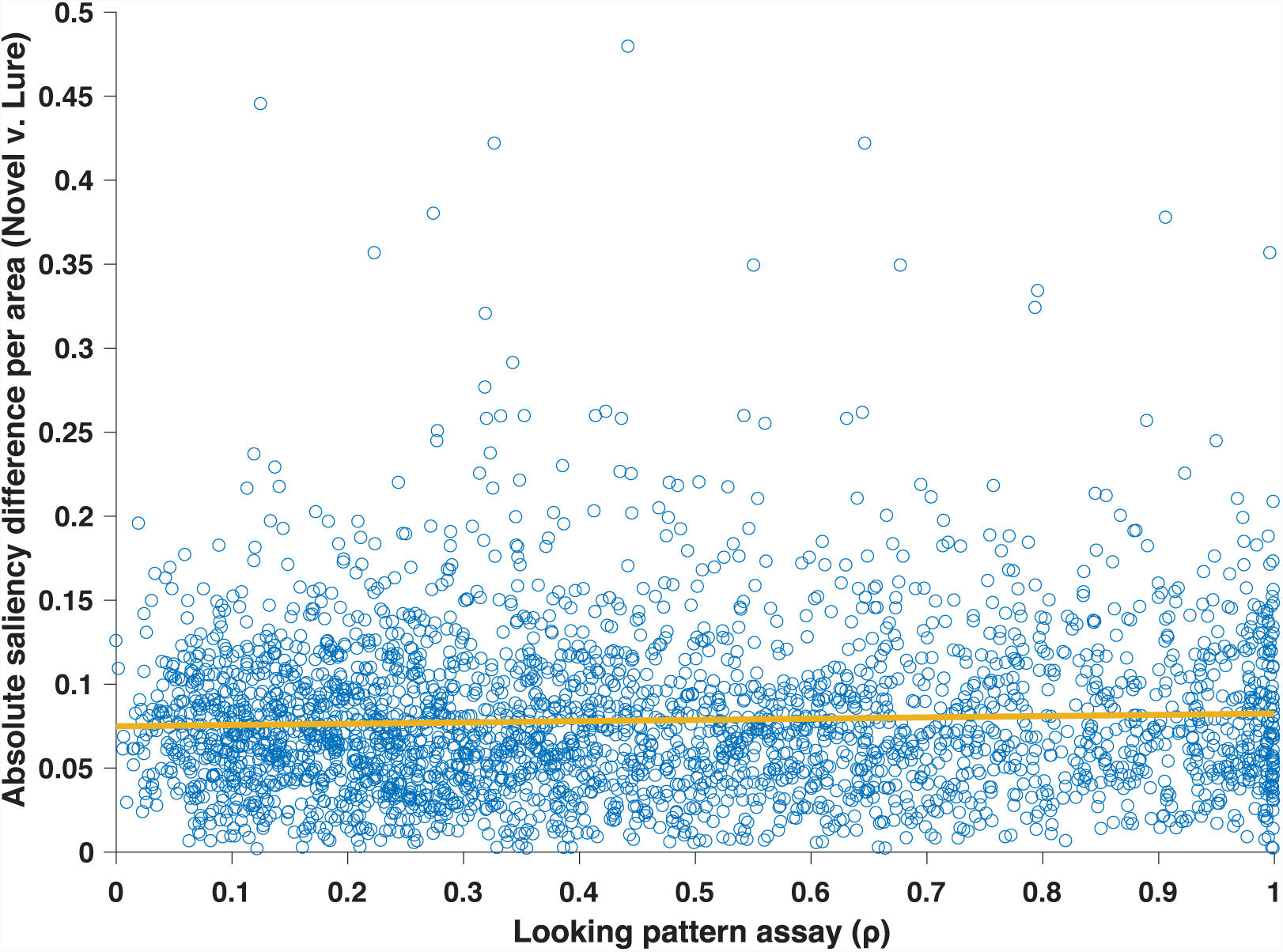
Saliency differences vs. looking pattern assay values. We correlated the absolute difference in saliency per area between each novel and lure image and then plotted that value versus the looking pattern assay value (**ρ**) for the monkey for each of the same pairs. If our looking pattern assay only measured the difference in salience between the novel and lure images, we would expect a negative relationship in this graph since low **ρ** values (indicative of differing scanpaths) would correlate with larger changes in salience. Instead, the trend is slightly positive, meaning the looking pattern assay is influenced by memory and not merely the perceptual differences between the novel and lure images.

**Figure S5:**
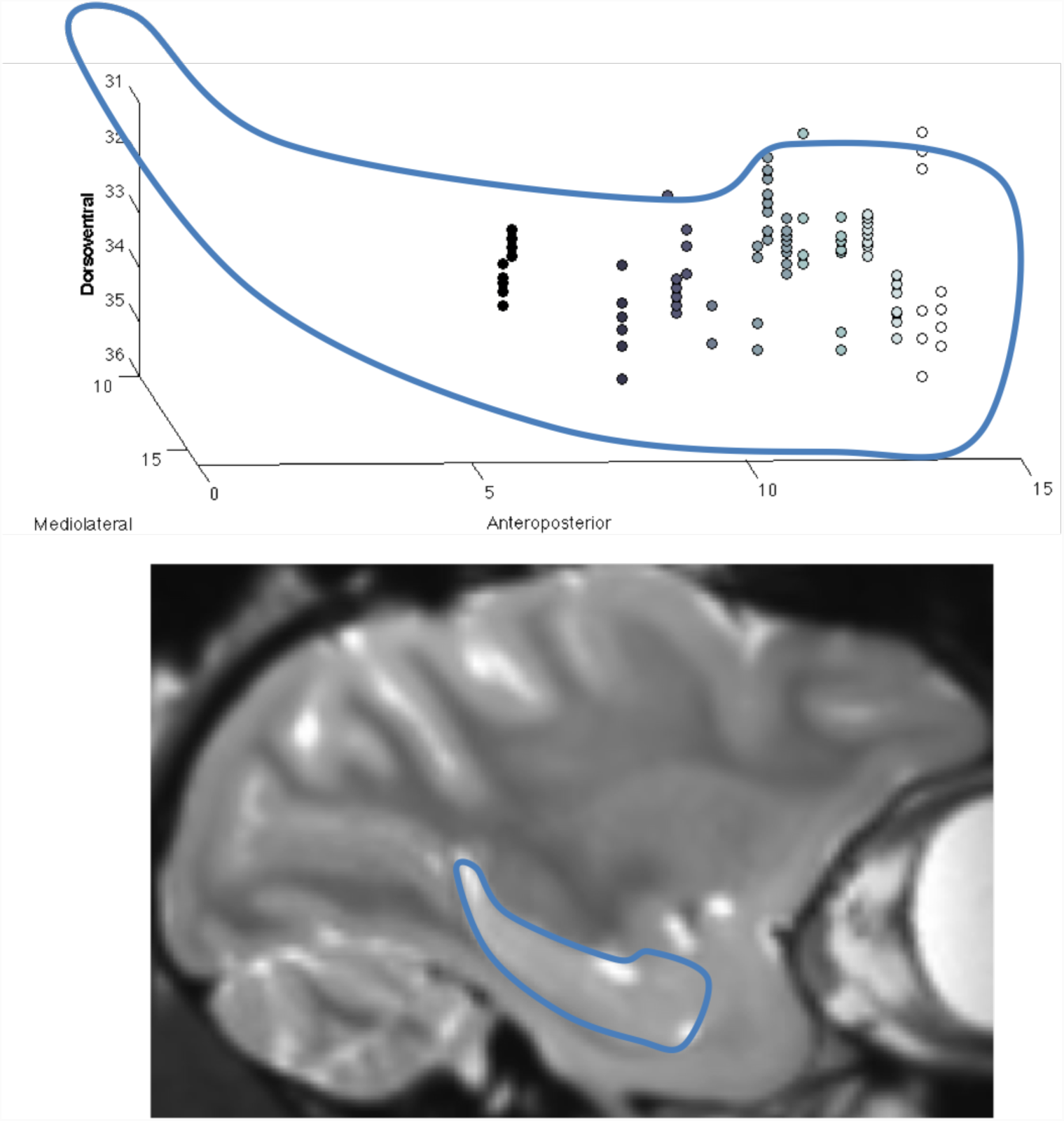
Recording sites superimposed on structural MRI of hippocampus. (Top) 3-dimensional view of recording locations from monkey I. Left to right shows the posterior to anterior direction, with colors from black to white indicating 6 to 14 mm. anterior of ear bars. (Bottom) Same outline as in top figure superimposed over sagittal MRI slice 13 mm. right of midline.

**Figure S6:**
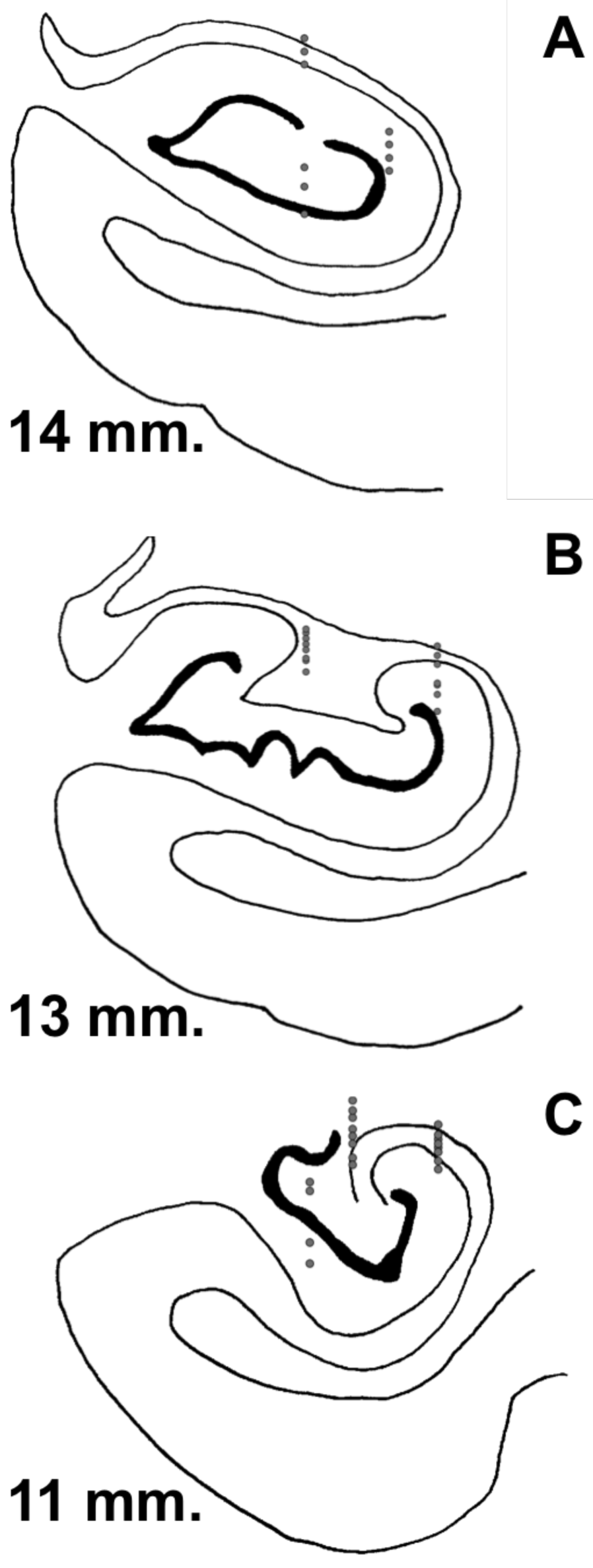
Recording sites superimposed on histology slices. Monkey I coronal slices at 3 anteroposterior levels of hippocampus with recording sites superimposed on estimated recording locations (dots). Slices are aligned in the mediolateral direction. Distance is relative to ear bars = 0 mm.

**Figure S7:**
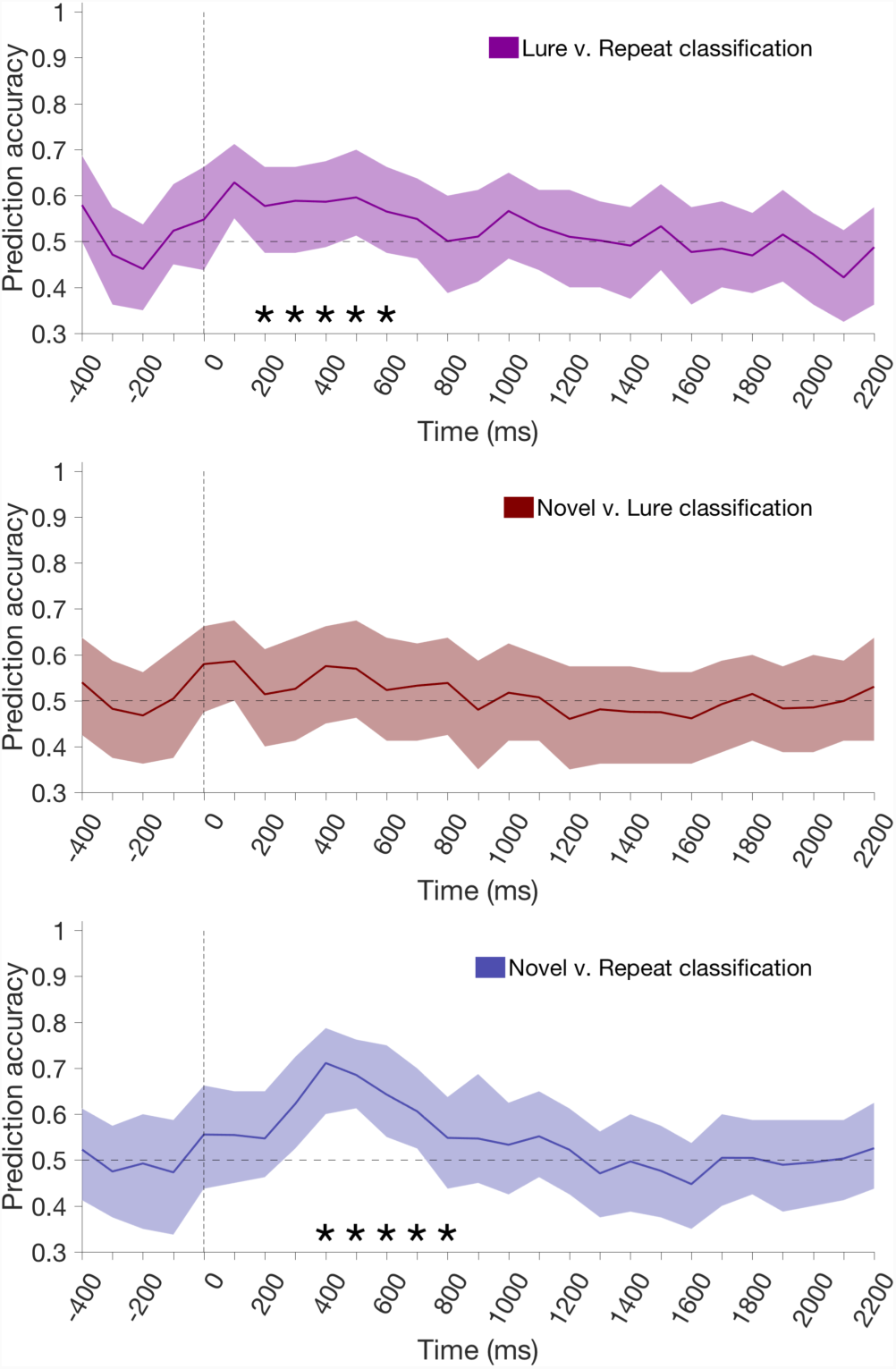
Prediction accuracy for classifier trained on neural population with only trials ≤1000 ms looking time. Same conventions and analysis as Fig. 2, except trials with monkey looking times outside the window of 300-1000 ms were excluded. Since this reduction lowered the trial count in our classifier, we lowered the minimum lure and repeat trials necessary for inclusion of a neuron in our population to be 40 instead of 50. This population came to 99 of the 110 neurons used in Fig. 2 for lure v. repeat classification.

**Figure S8:**
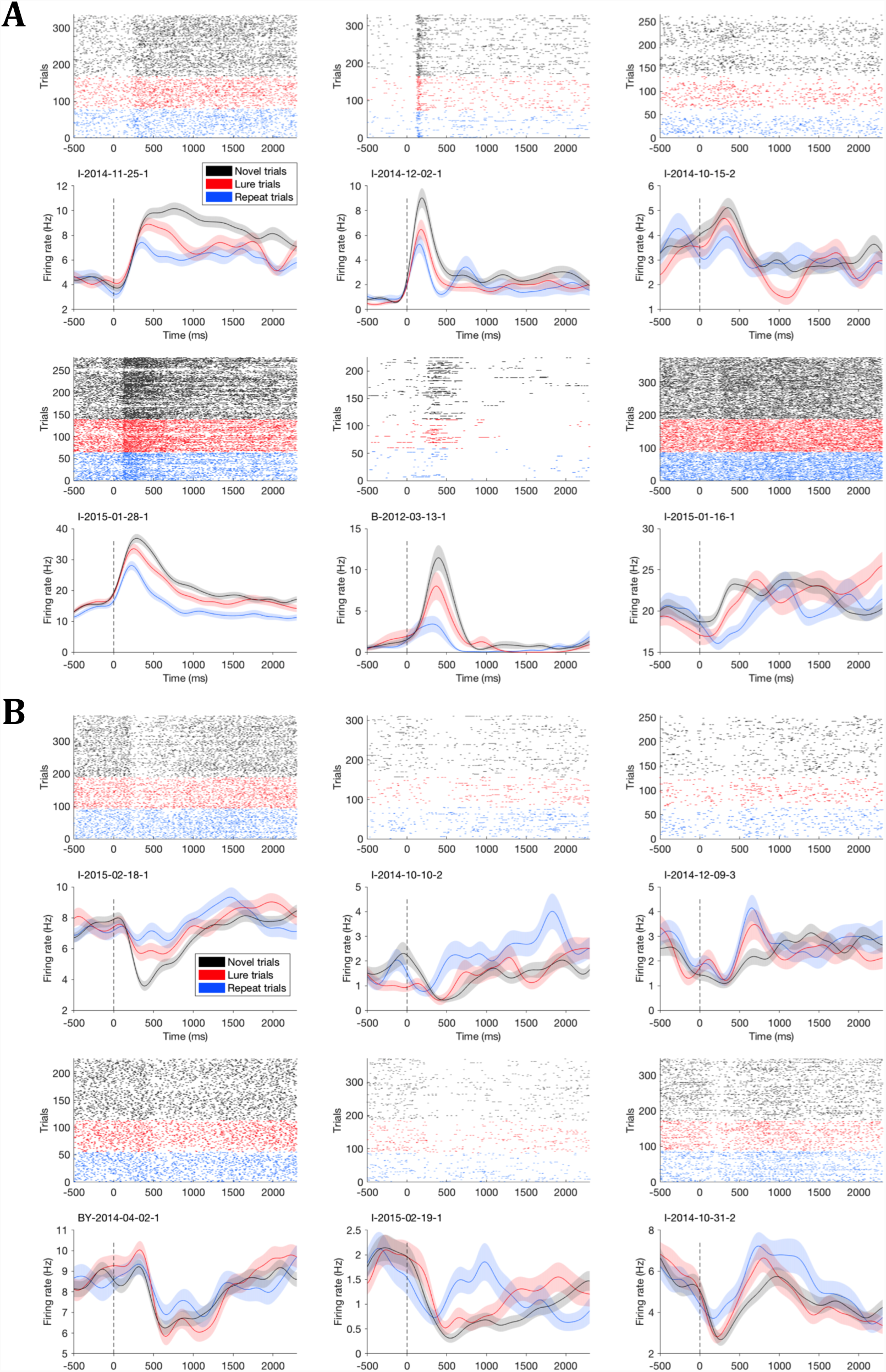
Raster plots of neurons that contribute most to image classification. Same as Fig. 3, but with raster plots above each spike density function. There are always as many novel trials as lure and repeats combined, since only novel trials that are the first presentation of the similar image as the lure and the first presentation of the repeat are included in order to keep image content matched.

**Figure S9:**
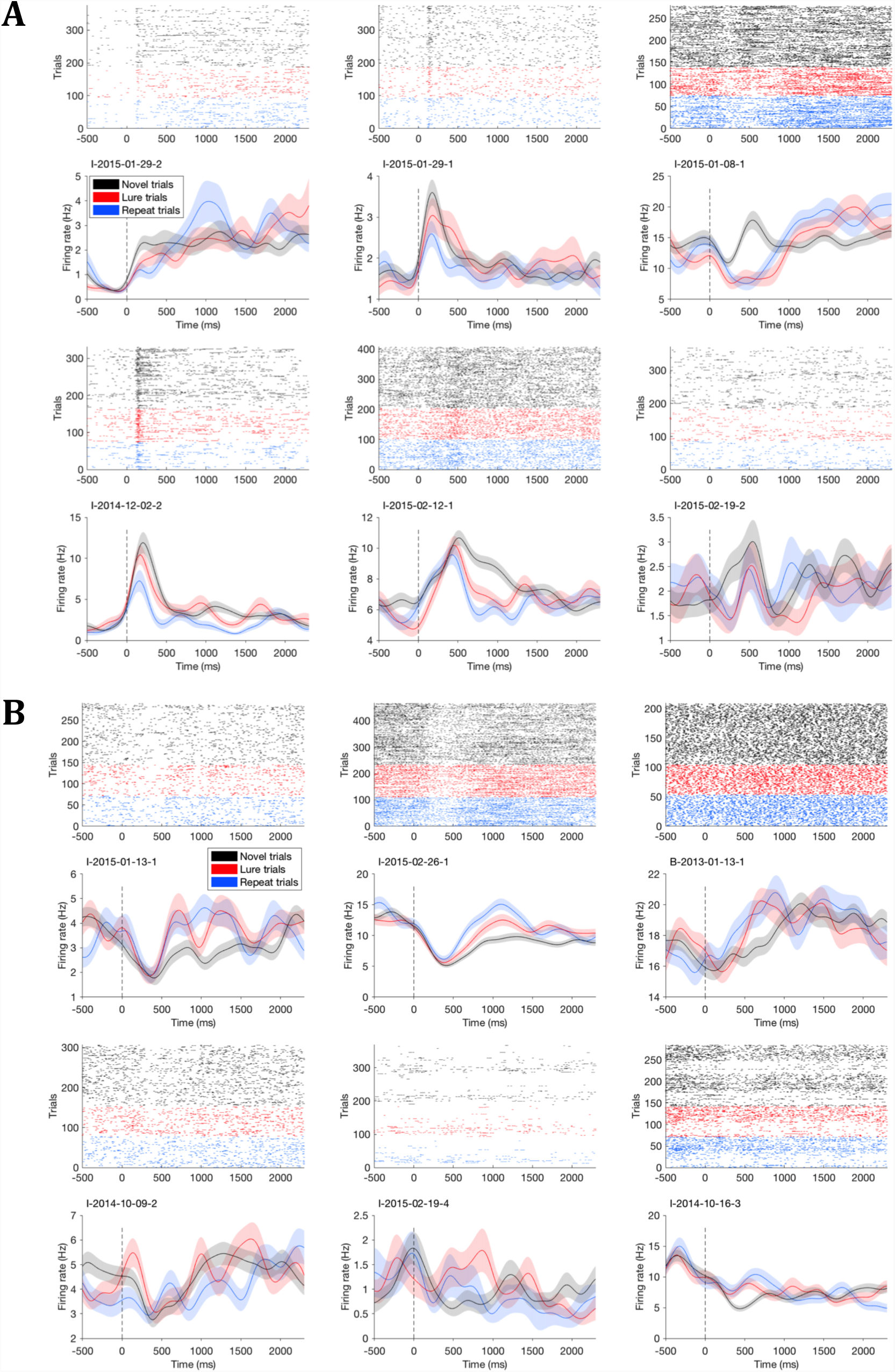
Raster plots of next 6 neurons that contribute most to image classification. Same conventions as Fig. S8, for the 7-12^th^ neurons with the most positive classifier weights **(A),** and the 7-12^th^ neurons with the most negative classifier weights **(B)**.

**Figure S10:**
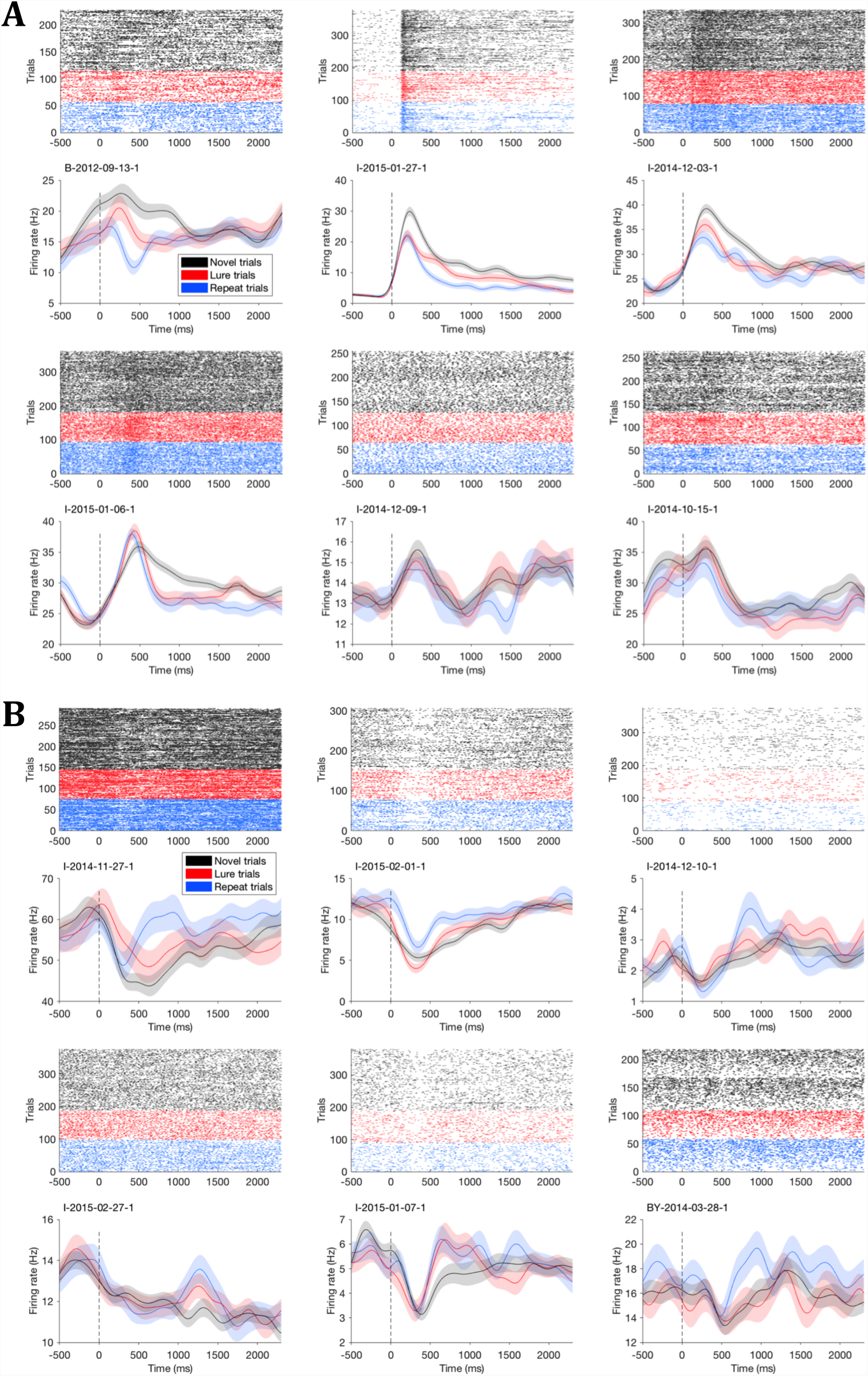
Raster plots of 6 additional neurons from each group. Same conventions as Fig. S8. Neurons were not selected by contribution to classifier, but **(A)** show qualitative similarities to positive regression weight group (i.e. early peaks with highest firing rates to novel images) and **(B)** qualitative similarities to negative regression weight group (i.e. early inhibition with highest firing rates to repeat images).

**Figure S11:**
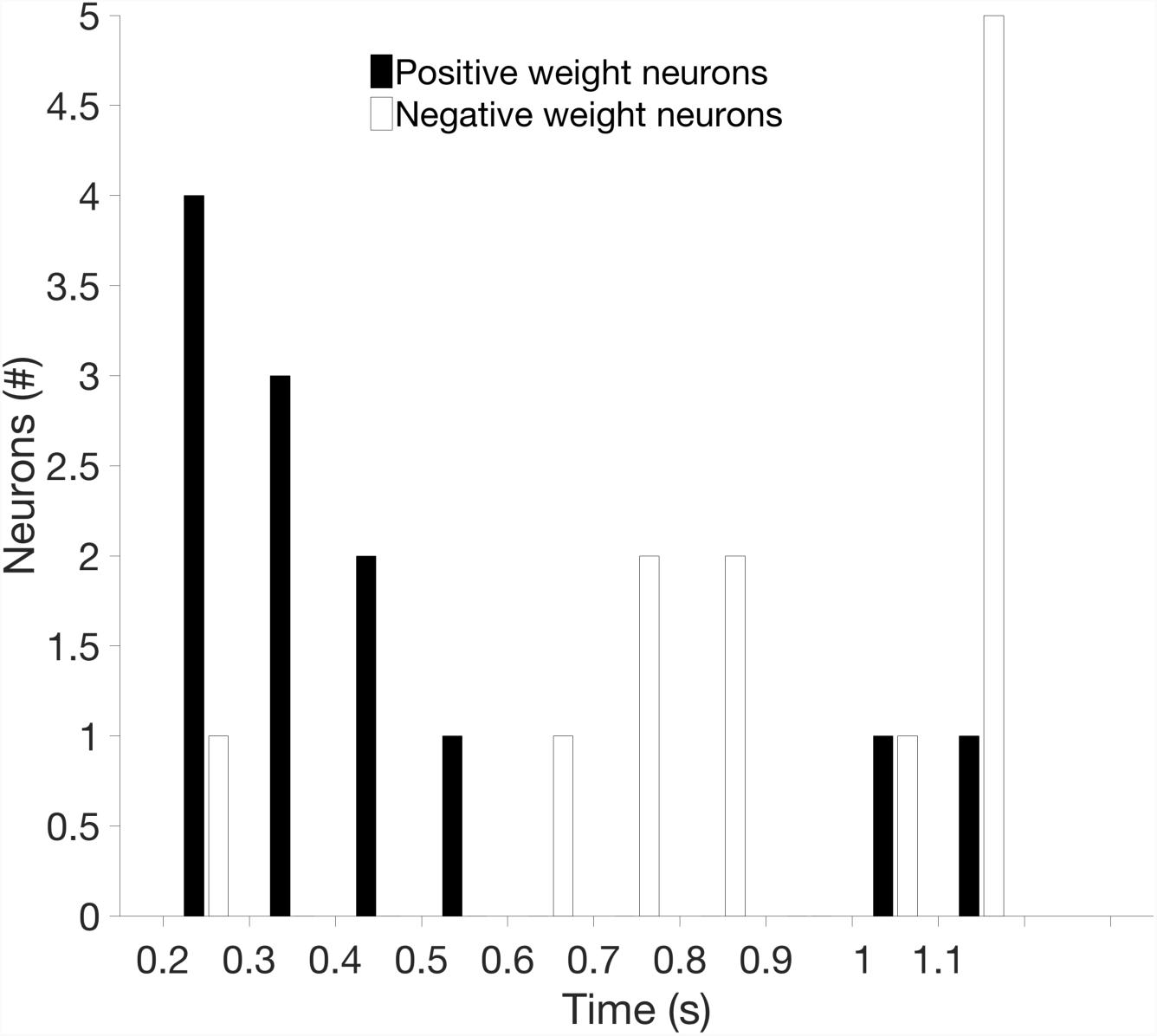
Histogram of peak firing rate time bin for positive and negative regression weight neurons. The time bin from 200-1200 ms with the peak firing rate averaged across all trials for each of the 12 neurons with the most positive regression weights (Figs. S8A-S9A) and the 12 neurons with the most negative regression weights (Figs. S8B-S9B).

**Figure S12:**
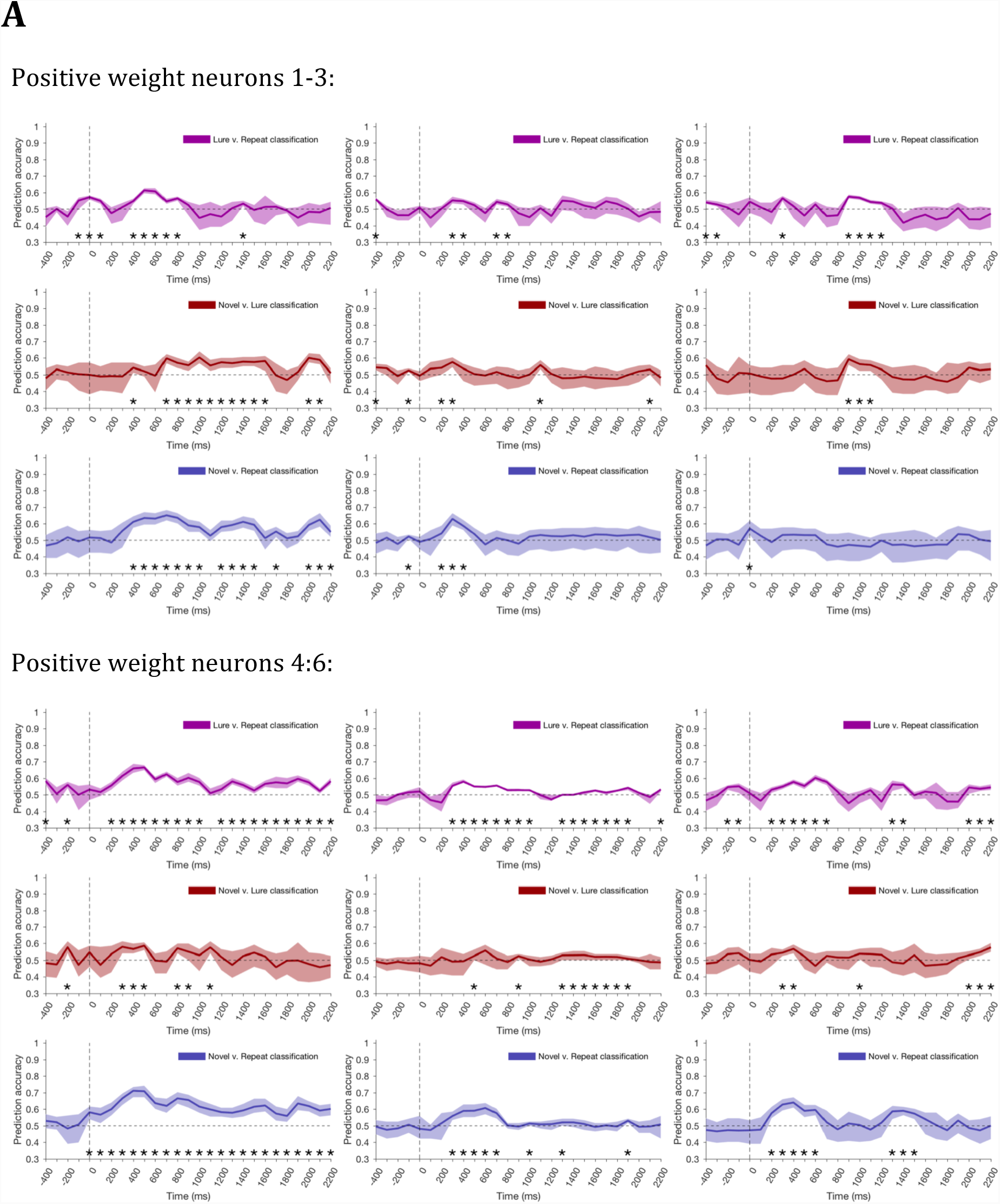
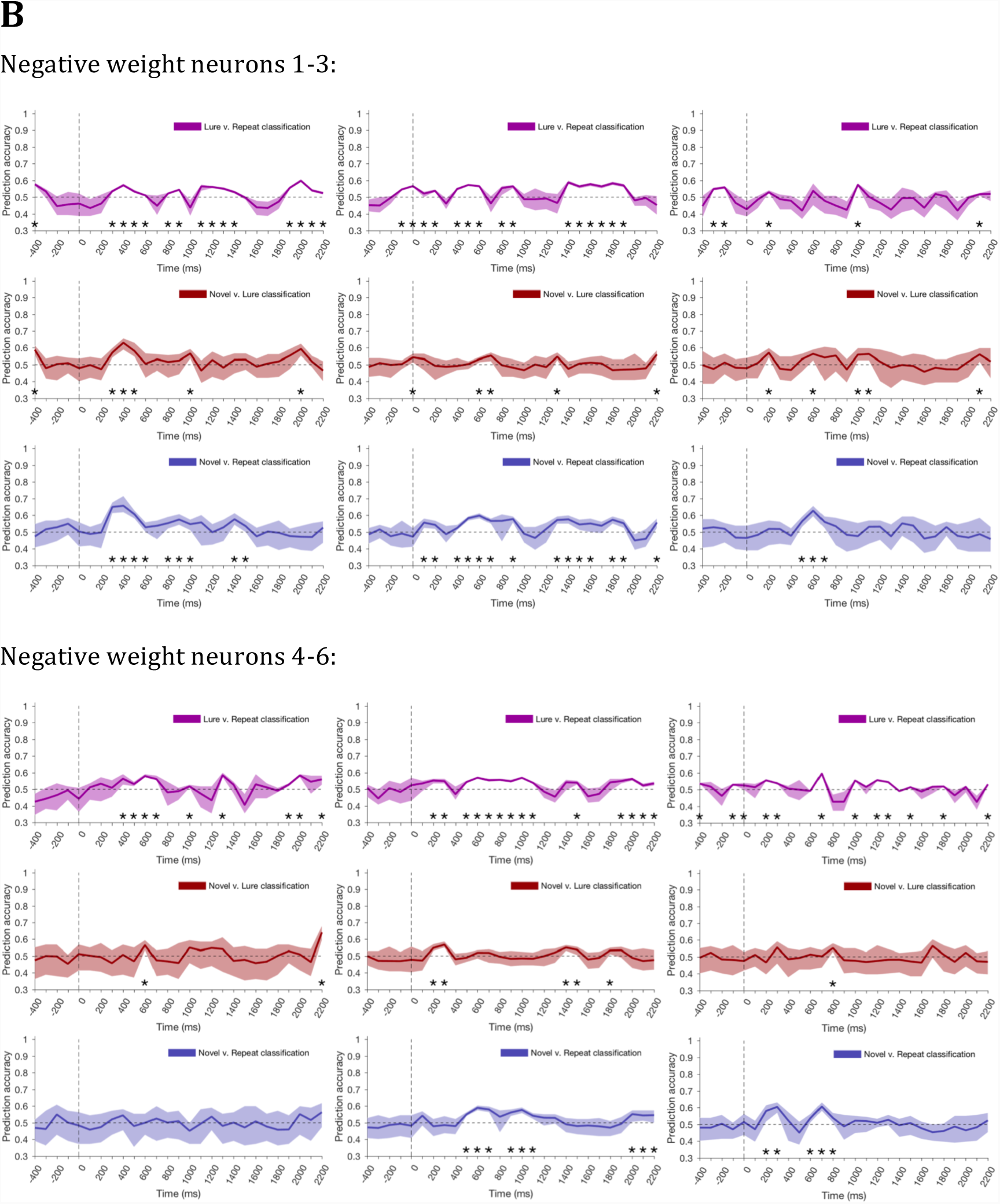
Prediction accuracy for classifier trained on single neurons. Similar to Figure 2, except logistic regression was used for the classifier instead of ridge regression. Each classifier was run through 100 permutations with subsampling of whichever trial type being compared had more trials. For example, if there were 75 lure trials and 80 repeat trials recorded for a single neuron, each permutation would use all 75 lure trials and randomly sample from the 80 repeat trials. As above, the prediction error was averaged across all permutations and error bars represent the 5^th^ and 95^th^ percentile accuracy scores at each separate time bin. **(A)** Same 6 neurons from Figs. 3A and S8A. **(B)** Same 6 neurons from Figs. 3B and S8B. Asterisks indicate significant accuracy as indicated by the 5^th^ percentile permutation eclipsing chance level classification (50%).

